# Capturing the interplay of membrane lipids and structural transitions in human ABCA7

**DOI:** 10.1101/2021.03.01.433448

**Authors:** Le Thi My Le, James R. Thompson, Sepehr Dehghani-Ghahnaviyeh, Shashank Pant, Phuoc X. Dang, Takahisa Kanikeyo, Emad Tajkhorshid, Amer Alam

## Abstract

Phospholipid extrusion by ABC subfamily A (ABCA) exporters is central to cellular physiology, although the specifics of the underlying substrate interactions and transport mechanisms remain poorly resolved at the molecular level. Here we report cryo-EM structures of lipid-embedded human ABCA7 in an open state and a nucleotide-bound, closed state at resolutions between 3.6-4.0 Å. The former reveals an ordered patch of bilayer lipids traversing the transmembrane domain (TMD), while the latter reveals a lipid-free, closed TMD with a small extracellular opening. These structures offer a structural framework for both substrate entry and exit from the ABCA7 TMD and highlight conserved rigid-body motions that underlie the associated conformational transitions. Combined with functional analysis and molecular dynamics (MD) simulations, our data also shed light on lipid partitioning into the ABCA7 TMD and localized membrane perturbations that underlie ABCA7 function and have broader implications for other ABCA family transporters.

## Introduction

ABCA family exporters mediate efflux of phospholipids and sterols from cells, contributing to membrane homeostasis, bilayer structure and asymmetry, and the formation of serum lipoproteins, among other key physiological processes^1^. Their dysfunction therefore underlies several human diseases^2–4^. The molecular details governing the ABCA exporter substrate transport cycle are not fully resolved. To fill this knowledge gap, here we present the structural and functional analysis of human ABCA7, whose dysfunction has been strongly linked to Alzheimer’s Disease (AD)^5–12^, in a lipid environment. Deficient ABCA7 activity leads to alterations in both brain lipid profiles and fatty acid and phospholipid biosynthetic pathways^13^, impaired memory, and reduced immune responses^14,15^. Both *in vitro* lipid flipping^16^ and lipid extrusion to apolipoproteins by cells over-expressing ABCA7^17^ have been demonstrated, although the correlation between the two processes, if any, remains unclear. To date, no direct structural information exists for ABCA7. Understanding the molecular details of the ABCA7 transport cycle and how its dysfunction alters inflammatory and immune responses, lipid homeostasis, and phagocytosis, which all contribute to AD progression^18–22^, may therefore pave the way for novel therapeutics for AD.

ABCA7 encodes a 2146 amino acid membrane transporter found in many tissues and blood, hippocampal neurons, macrophages, and microglia^23,24^. Like the phospholipid and sterol exporter ABCA1 and retinal importer ABCA4, with which it shares 54% and 59% sequence similarity, respectively, ABCA7 comprises two halves assembled as a full transporter. Each half consists of a TMD, with the first two transmembrane helices (TMs) of each separated by a large extracellular domain (ECD), and a nucleotide binding domain (NBD) attached to a cytoplasmic regulatory domain (RD). To visualize its ATP-dependent conformational cycle in a lipid environment, we resolved the structures of human ABCA7 in multiple conformations in lipid and detergent environments using cryo-EM and probed its lipid interactions using ATPase assays and MD simulations. Our data allow us to directly visualize lipid partitioning into the TMDs and the associated conformational changes in ABCA7 that provide insights into its mechanisms of substrate entry and export that likely hold true for other members of the ABCA family.

## Results

### Dependence of ABCA7 ATPase activity on its lipid environment

Human ABCA7 expressed in a tetracycline inducible stable HEK293 cell line was reconstituted in liposomes and nanodiscs comprising 80% brain polar lipids (BPL) and 20% cholesterol (Chol) and its ATPase activity was measured (Figure 1A, Figure S1A). Although ATP hydrolysis was slowest in nanodiscs, it followed Michaelis-Menten kinetics similar to ABCA7 in detergent or liposomes comprising the same lipid/cholesterol composition and Michaelis constant (K_M_) values for all three were in the 0.5-0.8 mM range. ATPase rates for a hydrolysis-deficient mutant carrying E965Q and E1951Q substitutions (ABCA7_EQ_) were drastically reduced compared to wildtype in both nanodiscs and detergents, demonstrating that the observed activity was specific (Figure S1A).

**Figure 1.**
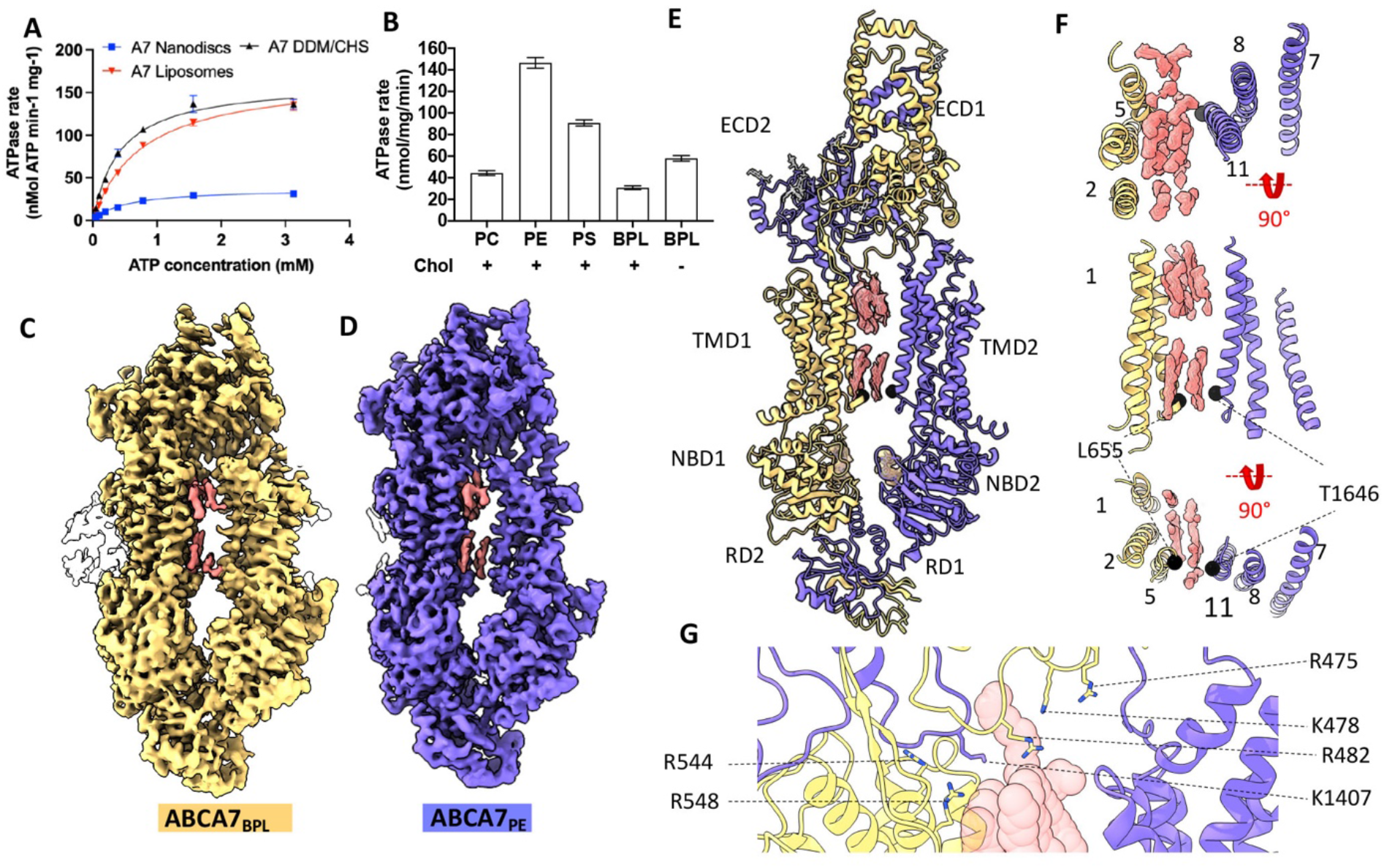
Functional characterization and structure of human ABCA7 incorporated in nanodiscs. **(A)** ATPase activity of ABCA7 at different ATP concentrations and **(B)** different nanodisc phospholipid/cholesterol compositions. Cholesterol (20%) presence is indicated by + or -. Experimental replicates (n)=3 and error bars represent standard deviation (s.d). (**C)** Cryo-EM map of ABCA7_BPL_ (yellow) at 3.6 Å resolution and **(D**) ABCA7_PE_ at 4.0 Å resolution(purple). Density for protein is shown in yellow (0.025 contour) and that for modeled lipid acyl chains (0.035 contour) shown in pink (TMD luminal lipids) and white (peripherally associated lipids). **(E)** ABCA7_PE_ shown in ribbon format with Half 1 colored yellow and Half 2 purple, along with density for TMD lipids (pink 0.025 contour) and bound nucleotide (yellow 0.035 contour). Acyl chains and glycans are shown as pink and grey sticks, respectively. (**F)** TMD lumen with density for TMD lipids (pink 0.025 contour) viewed from the extracellular side (top), membrane plane (middle) and cytoplasmic side (bottom). Cα atoms for the cytoplasmic gate are shown as black spheres. (**G)** View of the TMD-ECD interface with select residues oriented towards lipids (transparent red spheres) shown.

To analyze the effect of different lipid headgroups on ATPase activity, we compared ATPase rates of ABCA7 in BPL/Chol nanodiscs to those in which BPL was replaced by either brain polar phosphatidylethanolamine (PE), phosphatidylserine (PS) or phosphatidylcholine (PC) as well 100% BPL nanodiscs. As seen in Figure 1B, ATPase rates were highest in PE and PS nanodiscs followed by 100% BPL nanodiscs lacking cholesterol and, lastly, PC nanodiscs that showed a marginal increase. These results largely match previous findings^16^, although differences exist in extent of ATPase rate stimulation that are likely due to the divergent lipid formulations used for reconstitution. Overall, they demonstrate that lipids modulate ABCA7 activity in a species dependent manner and that cholesterol has an inhibitory effect on ATPase activity of ABCA7.

### Cryo-EM structures of ABCA7 in nanodiscs reveal an asymmetric, open TMD

To visualize ABCA7 in a lipid environment, we determined its cryo-EM structures in BPL/Chol nanodiscs (ABCA7_BPL_) at 3.6 Å resolution (Figure 1C, Figure S1B-E). Despite the addition of the non-hydrolysable ATP analog adenosine-5′-o-(3-thio-triphosphate) (ATPγS), the transporter adopted an open conformation with separated NBDs and a wide open TMD pathway. We observed density features consistent with a patch of ordered lipids from both bilayer leaflets traversing the width of TMD as expanded upon below. A second, higher-resolution 3D class displayed more complete density for the ECD tunnel region but comparatively lower quality of EM density for TMD2-NBD2 and the entire RD, indicating greater conformational disorder, which was therefore not analyzed further (Figure S1D-E). Analysis of TMD-ECD interfaces in the ABCA7_BPL_ structures revealed more extensive contacts between the ECD and TMD1 (buried surface area (BSA) of 790 Å^2^) compared to TMD2 (330 Å^2^ BSA). Both TMD1 and TMD2 made contacts with the opposite ECD subunits, leading to a domain swapped arrangement. The TMD1-ECD2 interface (BSA 565 Å^2^) was significantly larger than that of TMD1-ECD1 (BSA 223 Å^2^) and both interfaces comprised an extensive network of polar and electrostatic interactions. The RD adopted a domain-swapped arrangement similar to that in ABCA4 structures^25^, with RD1 and RD2 associated with the opposite NBDs. To see whether structural changes could rationalize the enhanced ATPase activity observed in ABCA7_PE_ compared to ABCA7_BPL_, we also determined the cryo-EM structure of the former to 4.0 Å resolution (Figure 1D, Figure S2). Interestingly, both structures were nearly identical to each other (r.m.s.d 0.31Å), although only a single high resolution 3D class was resolved for ABCA7_PE_ (Figure S2). Compared to ABCA7_BPL_, density for TMD lipids was more homogenous, despite its comparatively lower resolution, as described below, indicating overall decreased conformational heterogeneity. For both ABCA7_BPL_ and ABCA7_BPE_, greater positional disorder was observed for TMD2-NBD2 compared to TMD1 and NBD1 as indicated by relatively weaker EM density quality (Figure S3).

### The ABCA7 TMD lumen is accessible to an ordered file of bilayer lipids

Our ABCA7_BPL/PE_ structures are distinguished from available structures of ABCA1 and ABCA4 (all resolved in a detergent environment) by the wider open TMD lumen that is almost completely occupied by lipids. The observed lipids are continuous with the surrounding membrane except at the cytoplasmic leaflet near residues L655 and T1646 from TM5 and TM11, respectively (Figure 1E-F). EM density for the modeled lipids is recognizable by gaps between the extracellular and cytoplasmic leaflets and the two acyl chains of a single file of phospholipids in the two lipid leaflets. Towards the extracellular end, ECD residues R475, K478, R482, K1407 and TMD1 residues R544 and R548 form a cluster of positively charged side chains oriented towards the luminal lipids (Figure 1G). Overall, the luminal lipids are more closely associated with TMD1, with residues from TM2, TM5, and TM11 within 5 Å of the modeled acyl chains compared to only residues from TM11 from TMD2.

To compare the architecture of ABCA7 in the presence or absence of lipids, we also determined its structure in the detergent digitonin (Figure 2A-B, Figure S4A-E). This ABCA7_DIGITONIN_ structure revealed a similar overall conformation to that seen in structures of ABCA1^26^ and ABCA4^25,27,28^ (Figure S4F) with a more complete map for the ECD compared to ABCA7_BPL/PE_. As expected, no EM density for TMD lipids was visible, and only isolated density features attributable to detergent molecules associated with TMD1 were observed, although only at a contour level below where much of the surrounding detergent micelle was visible (Figure 2A,C). This is in contrast to ABCA7_BPL/PE_ structures where TMD lipid density was much stronger than that of the bulk nanodisc (Figure 2D-E), further validating our assignment of bilayer lipids. Compared to structures in nanodiscs, the TMDs in this ABCA7_DIGITONIN_ structure were arranged more symmetrically with respect to each other with a narrower lumen. Interestingly, EM density for TMD2 was still weaker than that of TMD1, in contrast to structures of ABCA4^25,27,28^ and ABCA1^26^ (Figure S4D) indicating that a more mobile TMD2-NBD2 pair may be a unique feature of ABCA7.

**Figure 2.**
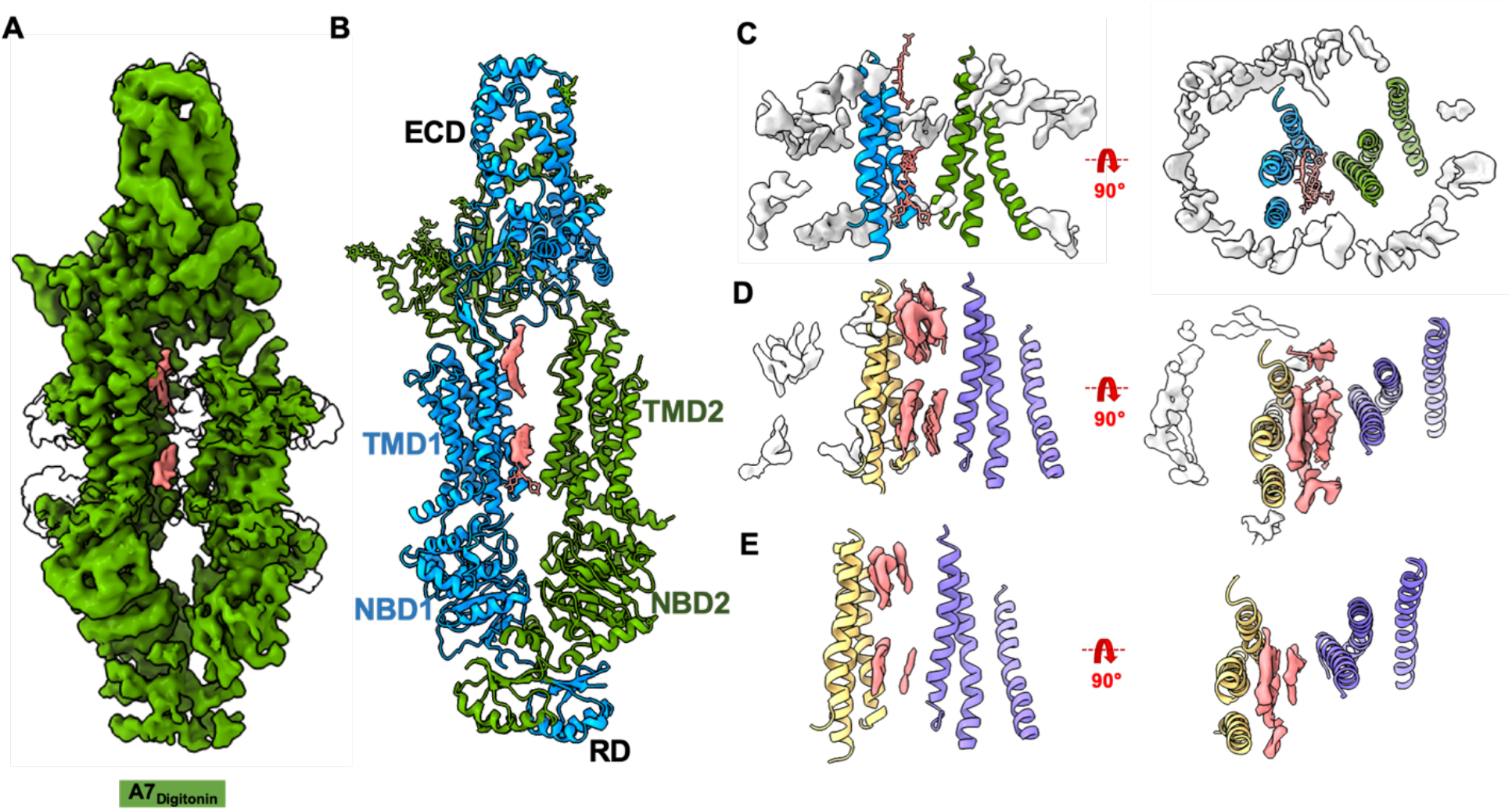
Cryo-EM structure of ABCA7_DIGITONIN_. **(A)** Cryo-EM map of human ABCA7_Digitonin_ at 3.9 Å with density for protein shown in green (0.013 contour) and extraneous density likely belonging to detergent shown in pink (0.013 contour). **(B)** Cryo-EM structure of human ABCA7_DIGITONIN_ shown in ribbon format with each half colored differently (blue and green). **(C)** TMD lumen of ABCA7_DIGITONIN_ with density for bulk micelle shown in white at 0.015 contour where detergent density inside TMDs disappears. Sticks for unmodeled detergent molecules are shown for which density is visible at the lower contour of 0.013. **(D)** TMD lumen of ABCA7_PE_ with density (0.035 contour) for TMD lipids (pink) and peripherally associated ordered lipids (white) shown. **(E)** Same as D with higher density contour of 0.046 where density of peripherally associated lipids is absent but that of luminal lipids remains.

### Closed TMD lumen and exit pocket for lipid extrusion in ATP bound ABCA7

To gain insight into possible mechanisms of lipid extrusion from the ABCA7 TMD, we used the hydrolysis-deficient ABCA7_EQ_ mutant reconstituted in BPL/Chol nanodiscs and determined its structure in a closed, ATP-bound state at 3.7 Å resolution (Figure 3A and Figure S5). This ABCA7_EQ-ATP_ structure revealed closely interacting TMD-NBD pairs and a largely occluded TMD lumen (Figure 3B). However, at the extracellular end of the TMD lumen, we observed a small central opening to the extracellular space/ECD that may represent an ‘exit pocket’ (Figure 3C,D) akin to that seen in the yeast pleiotropic drug resistance transporter Pdr5^29^ that could likely accommodate two acyl chains, (Figure 3D). TMD closure involves formation of a 4-TM bundle comprising TMs 2, 5, 8, and 11 that occludes the cytoplasmic bilayer leaflet. The exit pocket is comprised largely of hydrophobic residues, except for R548, which is part of a cluster of positively charged residues including R475, K478, R482 and R678 identified in our ABCA7_BPL/PE_ structures that may aid in directing a negatively charged phospholipid headgroup towards the ECD. Side chain density for this group of basic residues was poor compared to that in ABCA7_BPL/BPE_ structures, indicating greater disorder. TMD closure is accompanied by a rearrangement of both TMD-ECD interfaces compared to the open state structures (Figure S6). Compared to the ABCA7_BPL_ structure, both TMD1 and TMD2 interfaces with ECD1 increased from BSAs of 190 Å^2^ to 331 Å^2^ and from 76 Å^2^ to 285 Å^2^, respectively. Conversely, the BSAs of TMD1 and TMD2 with ECD2 decreased from 543 Å^2^ to 275 Å^2^ and from 317 Å^2^ to 192 Å^2^, respectively. As expected, the ABCA7_EQ-ATP_ structure shows a canonical NBD sandwich dimer with bound ATP (Figure 3E). In contrast to the ATP bound ABCA4 structure, RD density was too weak for accurate placement. However, contoured at lower thresholds, the available density features were more compatible with a rigid body shift rather than a conformational rearrangement of RD, with RD2 appearing to maintain contact with NBD1, but RD1 disengaged from NBD2.

**Figure 3.**
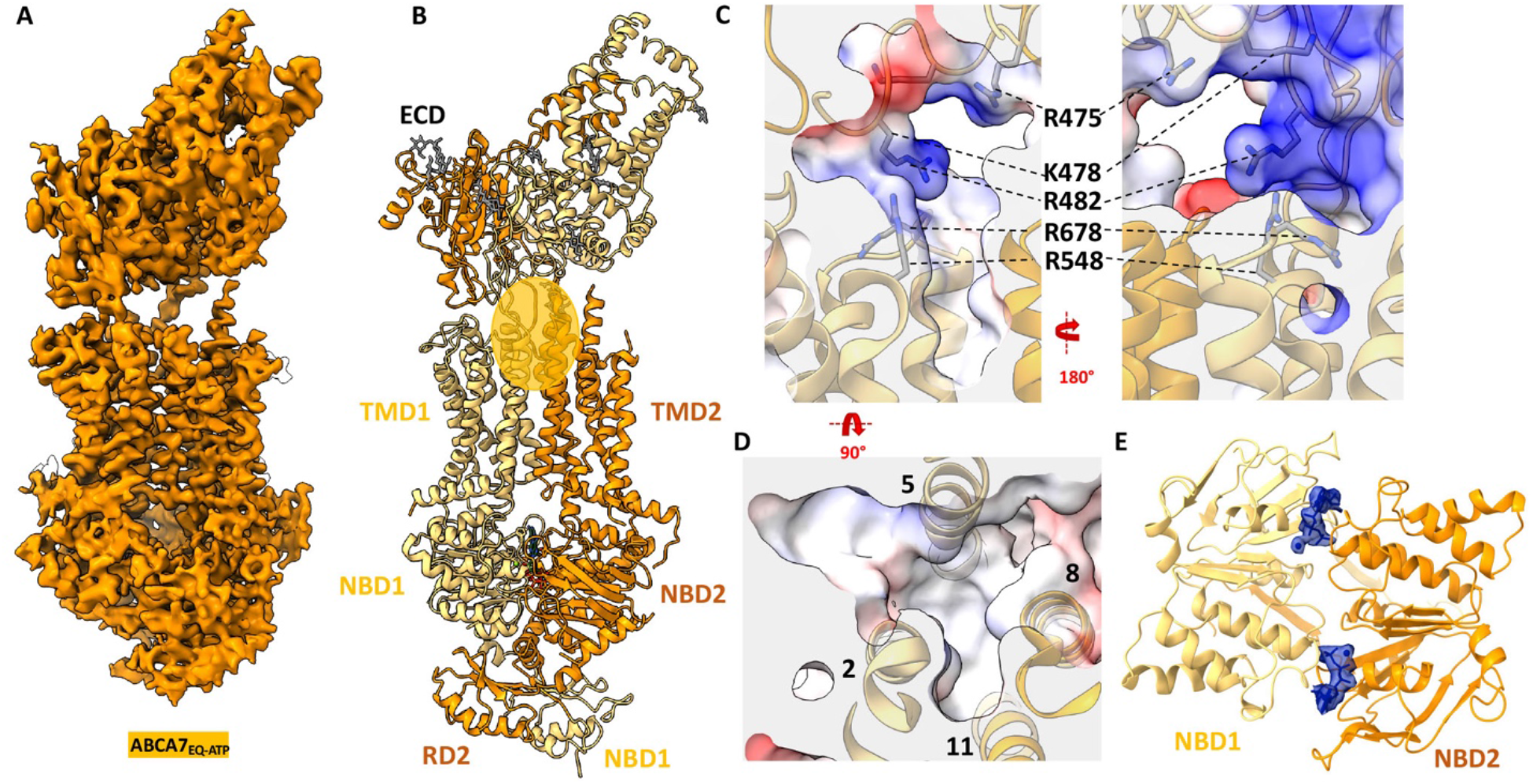
Cryo-EM structure of ABCA7_EQ-ATP_. **(A)** Cryo-EM map of human ABCA7_EQ-ATP_ in BPL/Chol nanodiscs at 3.7 Å resolution (orange density, 0.02 contour). **(B)** Cryo-EM structure of ABCA7_EQ-ATP_ shown in ribbon format with half 1 and half 2 colored yellow and orange, respectively. The transparent orange oval demarcates the observed exit pocket. **(C)** Central slice of electrostatic surface map showing surface details of exit pocket in two different orientations **(D)** 4-TM bundle forming the exit pocket. **(E)** NBDs viewed from the extracellular side with density for bound nucleotide (blue sticks) shown in blue (0.02 contour).

### Rigid-body motions of the TMDs define ABCA7 conformational transitions

Despite significant overall conformational differences, the individual TMDs in ABCA7_BPL/PE,_ ABCA7_DIGITONIN_, and ABCA7_EQ-ATP_ remain largely unaltered (Figure 4A). Moreover, the TMD-NBD pairs move as rigid body units from the fully open ABCA7_BPL_ structure to that of ABCA7_DIGITONIN_, which we assert is akin to an intermediate open state between the fully open ABCA7_BPL/PE_ state and the closed ABCA7_EQ-ATP_ states. The transition from open to intermediate open involves a rigid-body rotation of 9° and translation of 2 Å for TMD2-NBD2. The transition to the closed state involves a further 6° rotation and 15 Å translation of TMD2-NBD2 (Figure 4B-C), albeit with a greater alteration in the NBDs owing to a movement of their recA like domains upon ATP binding and dimerization. Rigid body movements were also observed for the entire RD in all three structures. The RD of ABCA7_DIGITONIN_ maintained molecular interactions with both NBD1 (RD2) and NBD2 (RD1), whereas the RD of ABCA7_BPL/PE_ separated from NBD2 while maintaining contact with NBD1. Finally, the ECD base region transitioned through a rigid body rotation of 34° and translation of 6.3 Å going from the open to closed states, whereas the tunnel and lid regions displayed greater heterogeneity.

**Figure 4.**
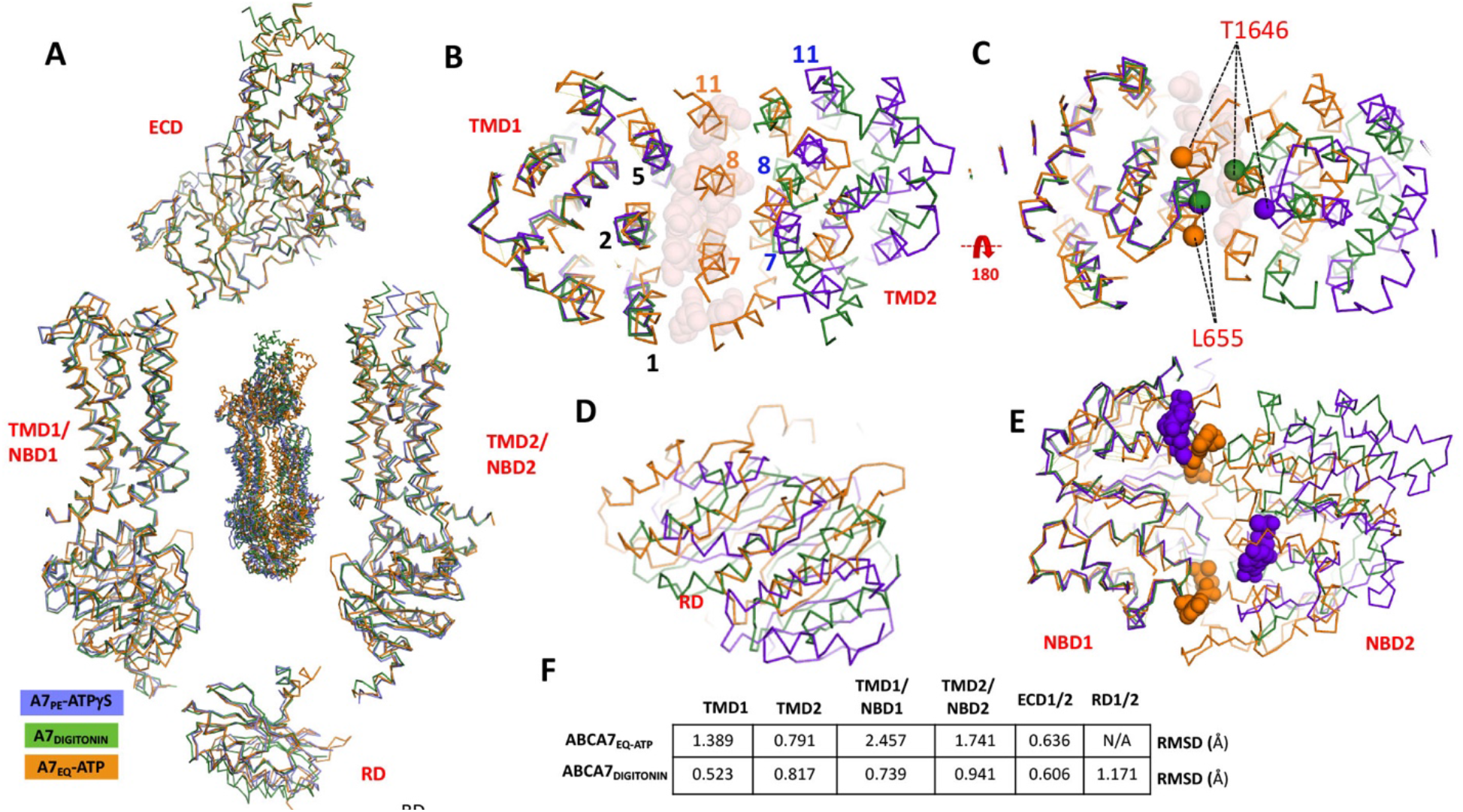
Comparison of open and closed conformations of ABCA7. **(A)** Overall structural alignment of the three ABCA7 conformations (center) along with individual alignments of rigid body pairs TMD1-NBD1, TMD2-NBD2, ECD, and RD (**B**) Overall alignment of the three ABCA7 conformations showing only TMD1 and TMD2 viewed from the extracellular side using the TMD1-NBD1 pair as an alignment reference. TMs lining the TMD pathway are numbered (**C**) Same as panel B, viewed from the cytoplasmic side with Cα atoms of gate forming residues shown as spheres (**D)** Same as panels B/C showing only RDs viewed from the cytosolic side. **(E)** Same as panel D showing just NBDs and bound nucleotides viewed from the extracellular side. ATPγS and ATP-Mg^2+^ are shown as purple and orange spheres, respectively. (**F)** Root mean square deviations (RMSD) of aligned atoms of ABCA7_BPE_ vs ABCA7_EQ-ATP_ and ABCA7_DIGITONIN_.

To establish whether the concerted TMD-NBD rigid body motions outlined above may extend to other ABC transporters of the Type-V ABC transporter/Type II ABC exporter fold^30^, we extended our analysis of individual TMD-NBD pairs to members of the G family for which the first structures of open and closed states were available, namely ABCG2 and ABCG5/G8 (Figure S7A)^31,32^. As shown in Figure S7B, the TMD-NBD pairs from both the apo and ATP-bound closed states of ABCG2, as well as from both halves of ABCG5/G8, all shared a similar overall architecture. Despite the divergent topologies of ABCG and ABCA transporters, with the former are arranged in an NBD-TMD configuration compared to TMD-NBD for the latter, the individual TMD-NBD pairs from ABCA7 shared very strong structural similarities with those of ABCG2, further establishing the role of rigid body movements of the TMDs to affect large scale conformational changes in these transporters.

### Lipid partitioning in the ABCA7 TMD captured with MD simulations

To gain additional molecular insight into the TMD-mediated lipid partitioning, membrane perturbation, and lipid extrusion from ABCA7, we performed multi-microsecond MD simulations of the open conformation of ABCA7_PE_ after embedding into two distinct lipid bilayers, one containing PE/Chol (4:1) and the other PC/Chol (4:1) (Figure 5, Figure S8). Each simulation was performed for 2 μs using a system including four copies of the protein (Figure 5A), resulting in an aggregate 8 μs sampling of lipid-protein interactions for each lipid composition. Interestingly, during the simulations, we observed PE and PC phospholipids penetrating the TMD cavity in both the cytoplasmic and extracellular leaflets in all four protein replicas (Figure 5A, insets). We quantified the number of phospholipids within the TMD cavity by counting the lipid headgroups (Figure S8C), revealing that on average, ABCA7 accommodated a larger number of PE lipids compared to PC (Figure 5B). To further quantify ABCA7-mediated deformation of the lipid bilayer bridging the TMDs, the average heights of phospholipid headgroups were calculated with respect to the midplane of the bilayer. We captured both a tendency for lipids to move towards the cytoplasmic gate and the formation of a dome-shaped phospholipid arrangement within the TMD lumen (Figure 5C-E, Figure S8A-B). Interestingly, our MD simulations showed an accumulation of phospholipid headgroups in the vicinity of residues R475, K478, R482, R544, and R548, with R482 and R548 displaying the most frequent lipid contacts (Figure S8D-E), corroborating our structural observations. Analysis of the cytoplasmic leaflet highlighted a continuous distribution of phospholipids with the surrounding membrane, except near TM5 and TM11 around residues L655 and T1646 (Figure 5D-E), which correlated well with the bilayer-like density in the ABCA7_BPL_ and ABCA7_PE_ cryo-EM maps (Figure 1C-D) that we modeled with acyl chains.

**Figure 5.**
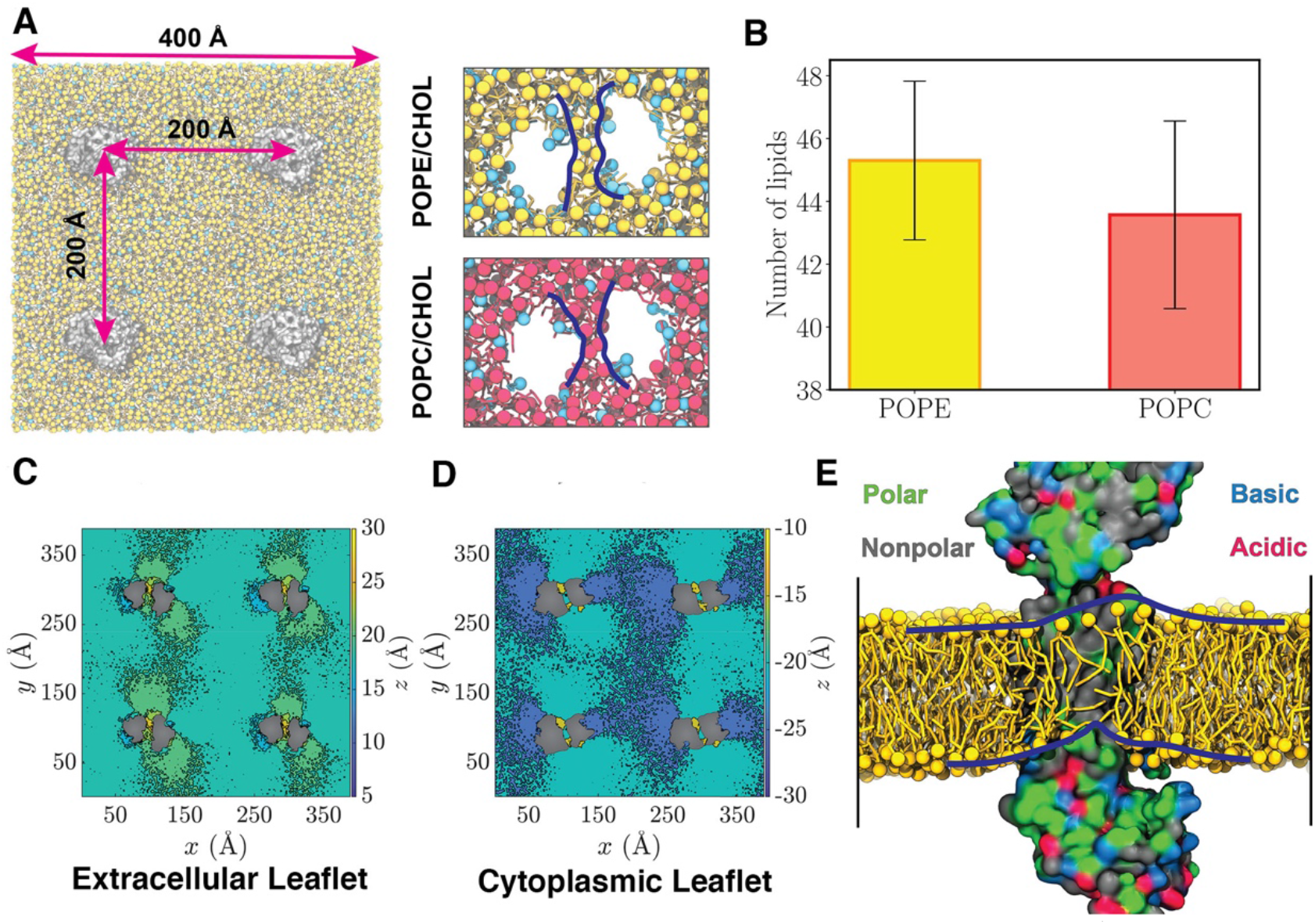
MD Simulations of ABCA7 in Lipid Bilayers. (A) Left: A representative simulation system with four copies of ABCA7 (silver), taken from the POPE/cholesterol (yellow/cyan) lipid patch. Right: the phospholipid belt (blue lines) formed in (top) the POPE/cholesterol (yellow/cyan) and (bottom) POPC/cholesterol (red/cyan) membranes (t = 2 μs). (B) The total count of phospholipids (sum of all four simulated proteins in a patch) partitioned in the TMD lumen for POPE/cholesterol and POPC/cholesterol bilayers (averaged over time ± standard deviation). (C/D) Heatmaps representing the average height (z values) of POPE headgroup with respect to the membrane midplane in the extracellular (C) and cytoplasmic (D) leaflets. Phospholipids are observed to climb the protein and form a dome-like configuration in the TMD lumen. (E) Snapshot of lipids partitioned in the TMD lumen, taken from the POPE trajectory at t = 2 μs. Polar, nonpolar, basic, and acidic residues are colored green, gray, blue, and red, respectively. TMD2 is hidden for a clearer view of the luminal dome-like lipid configuration (outlined by blue lines).

## Discussion

A mechanistic model derived from our results is shown in Figure 6. We establish that the TMD lumen of ABCA7 is accessible to bilayer lipids in the open state (ABCA7_BPL/PE_), providing a basis for substrate entry from both the extracellular and cytoplasmic leaflets. These ordered lipids are akin to those observed in the TMD cavity of AcrB, where they have been suggested to be important for functional integrity^33^. We identify a network of positively charged residues at the extracellular periphery of TMD1 and the base of the ECD directed towards the TMD luminal lipids. Our MD simulations support a role for these residues in interacting with phospholipid headgroups, consistent with our structural data. In ABCA7, R475 mutations have been identified in AD^34^ patients, and mutation of R638 in ABCA1 (equivalent to R548 in ABCA7) is associated with reduced serum HDL^35^ (Figures S9 and S10). While further studies are needed to pinpoint the specific roles, if any of these residues, in ABCA7 function, they remain largely conserved in ABCA1, ABCA4, and ABCA7. The upward protrusion of both bilayer leaflets within the TMD lumen suggested by MD simulations may, upon TMD closure, aid in sequestering phospholipids in the identified exit pocket to be primed for extrusion towards the ECD. Membrane deformations similar to those highlighted here have also been observed in the signal peptidase complex^36^, and further evidence the influence membrane proteins have on local membrane bilayer structure. ABCA7 could thus play a role in changing its membrane lipid environment by flipping cytoplasmic leaflet lipids to the extracellular leaflet and also extruding them back out into the bulk bilayer environment and/or to apolipoproteins. The reported defects in the phagocytotic activity of cells that express dysfunctional ABCA7^37,38^ could be related to ABCA7’s influence on the asymmetric distribution of bilayer lipids^39^, as enrichment of PS, shown to be flipped by ABCA7^16^, at the extracellular surface is linked to phagocytosis and phagocytosis-associated proteins within the membrane^40–42^.

**Figure 6.**
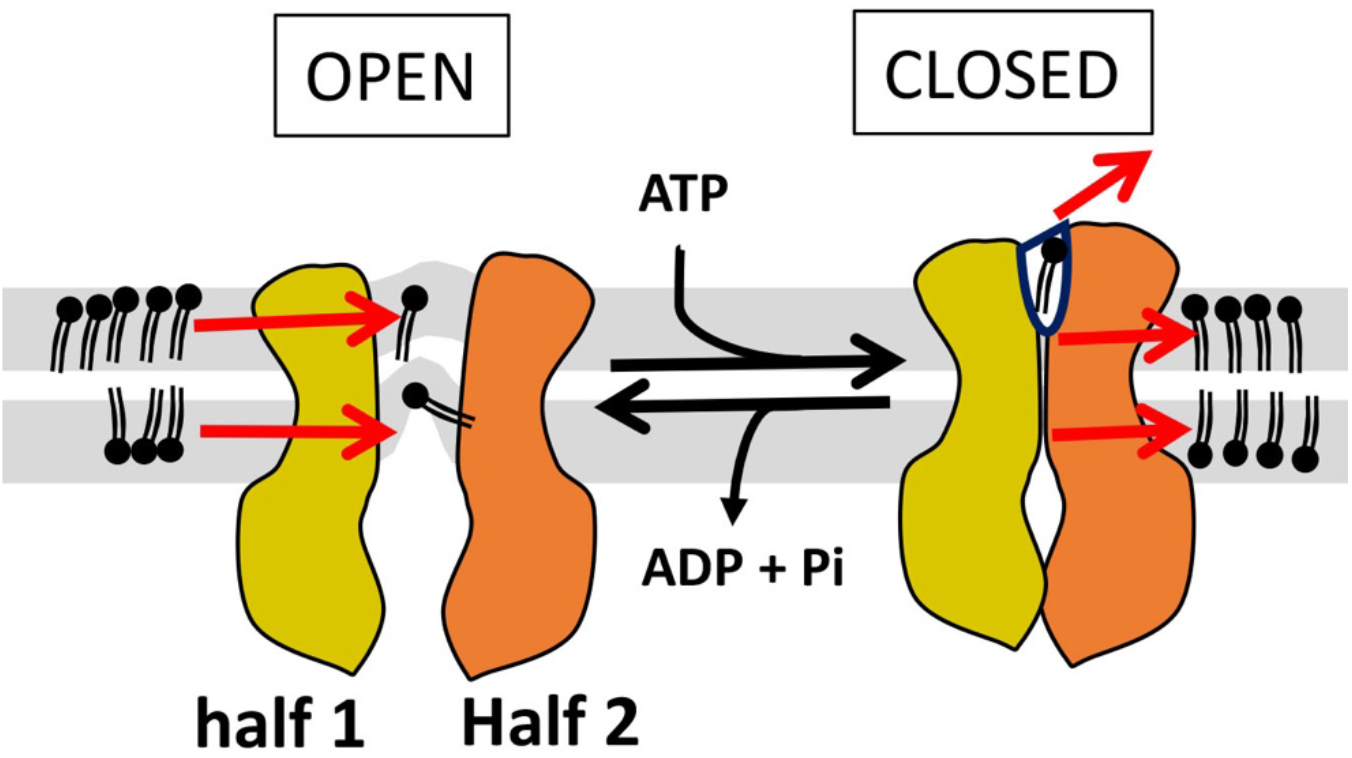
Proposed model of ABCA7 conformational transitions and lipid interactions. The membrane bilayer leaflets are shown in grey. Red arrows indicate movement of lipids (black) under the influence of TMD conformation. The ECD and RD have been omitted for simplicity as has any reference to potential lipid flipping from the cytoplasmic to extracellular leaflets.

Despite significant progress in our mechanistic understanding of ABCA transporter function, several questions remain open. First, it is unclear to what extent ABCA7 interacts with apolipoproteins in a physiologically relevant manner and, more broadly, the structural basis for apolipoprotein interactions with ABCA7 or ABCA1 is unknown. Second, while our structures provide a potential basis for lipid entry into and extrusion from the ABCA7 TMD, the mechanism whereby lipids get flipped, remains unresolved. Third, while our data point to a potential preference for PE over PC for TMD partitioning, the exact mechanism whereby ABCA1 or ABCA7 achieve lipid specificity are unknown. The observed enhancement in conformational homogeneity, quality of lipid density, and ATPase activity of our ABCA7_PE_ sample may be, in part, due to the smaller PE headgroup, which has been shown to aid in folding and stabilization of membrane proteins^43^. Finally, it is unknown how the opposite direction of substrate transport for ABCA4 is achieved considering similarities in ATP bound closed and open state structures of ABCA4 and ABCA7. Overall, our data will help devise better *in vitro* and *in silico* models to answer these questions, which will further aid in dissecting the unique roles these proteins play in cellular physiology.

## METHODS

### Protein Purification

We utilized the Flp-In TREX system (Thermo Fisher Scientific) for tetracycline inducible expression of human ABCA7. In short, a codon optimized synthetic gene construct (GeneArt/Thermo Fisher Scientific) of isoform 1 of ABCA7 (Uniprot ID Q8IZY2-1), harboring a C-terminal eYFP-Rho1D4 tag^43^ with a 3C/precision protease site between the protein and purification tags, was cloned into a PCDNA5.1 FRT/TO vector between BamHI and NotI restriction sites and a stable cell line was generated as per manufacturer’s protocol (Flp-In^TM^ T-Rex^TM^ Core Kit, Thermo Fisher Scientific). The resulting HEK293 based stable cells were grown and maintained in adherent cell culture in Dulbecco’s Modified Eagle Medium (DMEM, Thermo Fisher Scientific) supplemented with 9% Fetal Bovine Serum (FBS, Gibco) and a penicillin/streptomycin mixture (Thermo Fisher Scientific) at 37°C with 5% carbon dioxide (CO_2_) under humidified conditions. For protein production, cells were induced with 0.6 µg ml^-1^ tetracycline at a confluency of 80% in fresh DMEM supplemented with 2% FBS under otherwise identical conditions for an additional 72 hours before being washed with Phosphate Buffered Saline (PBS), harvested, and flash frozen in liquid nitrogen.

For purification, thawed cells were resuspended in a lysis buffer (Buffer L) comprising 25 mM Hepes pH 7.5, 150 mM sodium chloride (NaCl), 20% glycerol, 1 cOmplete EDTA free protease inhibitor tablet (Roche) per 50 ml Buffer L, 800 μM phenylmethylsulfonyl fluoride (PMSF) and 20 μg ml^-1^ soybean trypsin inhibitor (both Sigma), and mechanically cracked using a dounce homogenizer before addition of a 0.5%/0.1% w:v mixture of dodecyl maltoside (DDM) and cholesteryl hemisuccinate (CHS) (both Anatrace). Protein extraction was allowed to proceed for 90 minutes at 4°C with gentle agitation, after which, the suspension was centrifuged at 48,000 r.c.f for 30 minutes and the supernatant applied to rho-1D4 antibody (University of British Columbia) coupled Sepharose resin (Cytiva). Binding was allowed to proceed for 3 hours before the unbound fraction was discarded and beads rinsed with 4 x 10 bed volumes (BVs) of wash buffer (25 mM Hepes pH 7.5, 150 mM NaCl, 20% glycerol, 0.02%/0.004% w:v DDM/CHS). Protein was eluted by incubation with 3 BVs elution buffer (wash buffer supplemented with either 3C protease (1:10 w:w 3C:ABCA7) or 0.5 mg ml^-1^ 1D4 peptide (GenScript)) for 2-18 hours.

The EQ variant of ABCA7 contained two site mutations, E965Q and E1951Q. The E965Q site was created within the ABCA7 construct using site directed mutagenesis by PCR with the primers: A7eq1for 5’-GGTCATCCTGGATCAACCTACAGCAGGCGTGG-3’ and A7eq1rev 5’-GCCTGCTGTAGGTTGATCCAGGATGACCACC-3’. E1951Q was generated using a synthesized dsDNA block and the enzymes NheI and BsiWI. ABCA7_EQ_ with C-terminal eYFP-Rho1D4 tag was then transferred to a pCAG vector using KpnI and NotI restriction sites. The HEK293T cells were grown in Dulbecco’s Modified Eagle Medium (DMEM, Scientific) supplemented with 9% FBS (Gibco), penicillin/streptomycin mixture (Scientific) and antimycotic (Gibco) at 37°C and 5% CO_2_ under humidified conditions. A mixture of 37.5 µg of ABCA7_EQ_ plasmid and 75 µg of polyethyleneimine (PEI, Sigma) was incubated for 15 min at room temperature before being applied to 15 ml plates of HEK293T cells at a confluency of 60 - 80% to initiate transfection and expression. The cells were further cultured for 72 hours before being washed with PBS, harvested and flash frozen in liquid nitrogen. ABCA7_EQ_ was purified in the same approach of ABCA7; and eluted by incubation with 3 BVs elution buffer with either 3C protease for ATPase assay or with 0.25 mg ml^-1^ of 1D4 peptide in an additional 2 mM ATP and 10 mM MgCl_2_ for nanodisc reconstitution.

Membrane scaffold protein D1 (MSP1D1, addgene) and apoA1 were purified using established protocols for MSP^44^ with the following modifications: A synthetic construct of apoA1 bearing a 3C protease cleavable N-terminal deca-histidine tag (GeneArt/Thermo Fisher Scientific) was cloned into a pET28a vector (Addgene) and transformed in *E. coli* BL21 DE3 cells (New England Biolabs). One-liter cultures of Terrific Broth (TB) supplemented with 50 ug ml^-1^ kanamycin were grown from 10 ml overnight cultures from single colonies grown in LB. Cells were grown to an OD600 of 0.8 in a shaking incubator at 37 °C and induced with 1 mM isopropyl β-d-1-thiogalactopyranoside (IPTG). Protein expression was allowed to proceed at 20°C for 12 hours. Cells were centrifuged at 12,000 r.c.f, and pellets were flash frozen in liquid nitrogen and stored at -80°C until required. Frozen pellets were resuspended in 8 ml/ gram cell pellet resuspension buffer comprising 25 mM Hepes pH 7.5, 150 mM NaCl and 1 mM phenylmethylsulfonyl fluoride (PMSF) and sonicated. The suspension was spun down at 16,000 r.c.f at 4 °C for 30 min and the supernatant was applied 5 ml Ni-NTA resin (Qiagen)/ L culture medium. After discarding the flowthrough, the resin was washed with 25 mM Hepes pH 7.5, 150 mM NaCl, 1 mM PMSF and 20 mM imidazole until a pre-established baseline A280 reading was achieved. ApoA1 was eluted in 4 BVs of 25 mM Hepes pH 7.5, 150 mM NaCl, 1 mM PMSF and 200 mM imidazole, concentrated using a 10 kDa molecular weight cutoff (MWCO) Amicon filter (Millipore-Sigma) and desalted using a PD10 column (Cytiva) into 25 mM Hepes pH 7.5, 150 mM NaCl. The concentration of apoA1 was adjusted to 1 mg ml^-1^ for flash freezing in liquid nitrogen and storage at -80°C.

### ABCA7 and ABCA7_EQ_ nanodisc and proteoliposome preparation

For nanodisc reconstitution, peptide eluted or 3C cleaved ABCA7 was mixed with MSP1D1 and a mixture of BPL (brain polar lipid extract from Avanti) and cholesterol (80:20 w:w) with 0.5%/0.1% DDM:CHS using a 1:10:350 (ABCA7:MSPD1:lipid mix) molar ratio in nanodisc buffer (25 mM Hepes pH 7.5, 150 mM NaCl) that contained up to 4% glycerol for 30 minutes at room temperature (RT). Nanodisc reconstitution was induced by removing detergent with 0.8 mg ml^-1^ pre-washed Biobeads SM-2 (Bio-Rad) for 2 hours with gentle agitation at RT. For different phospholipid compositions, BPL was replaced with brain PE, PS, or PS (all from Avanti Polar Lipids). For structural studies, nanodisc-reconstituted ABCA7 bearing the eYFP-Rho1D4 tag was bound to rho-1D4 resin for an additional 2 hours, washed with 4 BV of nanodisc buffer, and eluted with 3C protease for 2 hours at 4°C. The eluted ABCA7 nanodiscs were concentrated using a 100,000 MWCO kDa Amicon filter and further purified by size exclusion chromatography using a G4000swxl column (TOSOH biosciences) equilibrated with nanodisc buffer at 4°C. Figure S1A shows a SEC chromatogram for pure ABCA7 placed into BPL/Ch nanodiscs, while a SEC chromatogram for ABCA7 in PE/Ch nanodiscs is in Figure S2A. Generally, three fractions were pooled from the main resultant peak.

ABCA7_EQ_ was reconstituted in nanodiscs for cryo-EM preparation using the same approach of ABCA7; except that an additional 2 mM ATP and 10 mM MgCl_2_ were present until the end of the purification procedure prior to grid preparation. A SEC chromatogram for ABCA7_EQ_ in BPL/Ch nanodiscs is shown in Figure S5A, where the trace is affected by the additional ATP added during the run.

ABCA7 proteoliposomes were generated by mixing detergent purified ABCA7 with liposomes at a protein: liposome ratio of 1:10 w:w. Liposomes were prepared by extruding a 20 mg ml^-1^, 80:20 w:w BPL/Ch lipid mixture 11 times using a previously described protocol ^45^. Briefly, detergent purified ABCA7 and liposomes were added to 0.14% and 0.3% Triton X100 (Sigma), respectively, then incubated for 30 minutes at RT. These two samples were mixed and incubated for 60 minutes. Detergent was removed by adding 40 mg fresh Biobeads SM2 (Bio-Rad) per ml reaction mixture during five successive incubation steps, 30 minutes at RT, 60 minutes at 4°C, overnight at 4°C, and two periods of 60 minutes at 4°C with gentle agitation. The suspension was centrifuged at 80,000 rpm for 20 minutes in an ultracentrifuge. The supernatant was removed, and the liposomal pellet was washed once with reconstitution buffer containing 150 mM NaCl, 25 mM Hepes pH 7.5. The ABCA7-liposome suspension was then centrifuged to remove the supernatant, and the proteoliposomes were resuspended at a final concentration of 0.5 - 1 mg ml^-1^ for ATPase assays.

### ABCA7_DIGITONIN_ preparation

For the digitonin solubilized ABCA7 purification, ABCA7 was extracted from the cell in a lysis buffer (Buffer L) using the same approach above, and the supernatant was applied to rho-1D4 resin for a 3-hour binding period. Then, the resin was rinsed with the 4 x 10 BVs of wash buffer containing 25 mM Hepes pH 7.5, 150 mM NaCl, 20% glycerol (v/v), and 0.06% digitonin (w/v). Protein was eluted by incubation with 3 BVs elution buffer, which was wash buffer supplemented with 3C protease (1:10 w:w 3C:ABCA7). Interestingly we obtained better particle distribution and ice quality with addition of a 1:2.5 molar excess of apoA1, prepared in house, to 3C cleaved ABCA7 prior to grid preparation. The mixture was concentrated by a 100,000 MWCO kDa Amicon filter (Millipore) and further purified by size exclusion chromatography using a G4000swxl column (TOSHOH biosciences) equilibrated with a buffer containing 25 mM Hepes pH 7.5, 150 mM NaCl, 0.035% digitonin (w/v), as shown in Figure S4A. Peak fractions were pooled and concentrated for cryo-EM grid preparation.

### ATPase assays

ATPase assays were based on a molybdate based colorimetric assay^46^. Protein concentrations used were in the range of 0.05-0.1 mg ml^-1^. Assays were started by the addition of either 2mM ATP, except for experiments in Figure 1B where 6.25mM ATP was used, in the presence of 10 mM magnesium chloride (MgCl_2_), incubated for 30 minutes at 37°C, then stopped by addition of 6% SDS. The assay was also performed in the presence of ABCA7 inhibitors as additives, such as 5 mM ATPγS or sodium orthovanadate. For ATP K_M_ measurements, a range of ATP concentrations was used. Statistical analysis was done using GraphPad Prism 9. ATPase rates were measured using simple linear regression, and the K_M_ of detergent, liposome, and nanodisc reconstituted ABCA7 were determined from the fit to the Michaelis-Menten equation of the corresponding rates. ABCA7 concentrations were measured using gel densitometry analyzed in ImageStudio Lite (LI-COR Biosciences) based on detergent purified ABCA7 standards with known concentrations determined by A280 measurements. All reaction components were mixed with ABCA7 in detergent or reconstituted in nanodiscs and liposome in the absence of ATP, incubated for 10 minutes at 37 °C prior to addition of ATP to start the reaction.

### Cryo-electron microscopy grid preparation

For ABCA7 reconstituted in nanodisc prepared in 80:20 w:w BPL/cholesterol and PE/cholesterol, SEC purified protein was mixed with 5 mM ATPγS (TOCRIS) and 5 mM MgCl_2_ for 20 minutes at room temperature and concentrated to 0.5 - 1.0 mg ml^-1^. Peak fractions from SEC already containing 2 mM ATP (Sigma) and 10 mM MgCl_2_ were pooled and concentrated to 0.5 - 1 mg ml^-1^ for nanodisc reconstituted ABCA7_EQ-ATP_. For ABCA7 in digitonin, peak fractions were pooled and concentrated between 2 to 5 mg ml^-1^. 4 µl samples were applied to glow discharged Quantifoil R1.2/1.3 grids (Electron Microscopy Sciences, Hatfield, PA, USA) using a Vitrobot Mark IV (Thermo Fisher Scientific) with a 4s blotting time and 0 blotting force under >90% humidity at 4°C, then plunge frozen into liquid ethane. For the nanodisc reconstituted ABCA7_EQ-ATP_ and ABCA7 in digitonin, two sample droplets were applied to glow discharged grids to obtain more particles per hole.

### Cryo-electron microscopy data collection and processing

Grids were clipped as per manufacturer guidelines and cryo-EM data was collected using a Titan Krios electron microscope operating at 300kV and equipped with a Falcon 3EC direct electron detector (Thermo Fisher Scientific.). Automated data collection was carried out using EPU 2.8.0.1256REL software package (Thermo Fisher Scientific) over multiple sessions in counting mode at a nominal magnification of 96,000x, corresponding to a calibrated pixel size of 0.895 Å for nanodisc reconstituted ABCA7_BPL_. Image stacks comprising 60 frames were collected at a defocus range of -0.6 to -2.6 μm and estimated dose rate of 1 electron/Å^2^/frame and further processed in Relion-3.1 (beta). Motion correction was done using Motioncor2 (Relion implementation)^47^ and contrast transfer function (CTF) correction was performed using Gctf 1.06^48^. A summary of the overall data processing scheme and the quality was presented in Figure S1C-E. In brief, 11802 micrographs were used for template free picking of 6725108 particles, followed by particle extraction at a 3x binned pixel size of 2.685 Å/pix. The dataset was processed in two batches. After 2-3 rounds of 2D classification 1259324 particles from Set 1 and 1088487 particles from Set 2 were selected for independent 3D classification steps (number of classes (K)=8 for both). The structure of human ABCA1 (EMDB6724) was used as a 3D reference for an initial 3D classification of a subset of the total data to yield an initial sub-nanometer resolution map of ABCA7 that was used as a 3D reference for the full datasets. After 1 round of 3D classification, both sets of data yielded a similar ensemble of classes. A total of 113291 particles from similar looking classes (black boxes) were subjected to an additional round of classification (K=3), ∼80% of which fell into a high-resolution class that yielded a 3.6 Å map after refinement and particle polishing steps. Similarly, 124114 particles from a second set of two similar classes (red boxes in Figure S1D) were selected for subsequent refinement, particle polishing, and post processing to yield a 3.1 Å map. All resolution estimates were based on the gold standard 0.143 cutoff criterion^49^. Datasets of ABCA7_PE_, ABCA7_EQ-ATP,_ and ABCA7_DIGITONIN_ were collected at a nominal magnification of 96,000x corresponding to a calibrated pixel size 0.889 Å, and image stacks containing 40 frames were collected a defocus range of -0.8 to -2.6 µm with an estimated dose rate of 1 electron/Å^2^/frame and further processed using the same software versions as for the ABCA7_BPL_ dataset unless otherwise indicated.

For ABCA7_PE_, a total of 2849251 particles were picked from 9218 ctf corrected (Gctf) micrographs in Relion ver. 3.1 in two batches at a 3-fold binned pixel size of 2.667 Å/pixel (Figure S2B-D). Of these, 783655 particles and 413330 particles were selected from Batch 1 and Batch 2, respectively and independently subjected to 3D classification (K=8) using a low pass filtered (60 Å) map of our ABCA7_BPL_ structure as a reference. A single highest resolution class from each was selected and their particles combined and subjected to an additional round of 2D classification. 2024649 particles were subjected to another round of 3D classification (K=8). A single, highest resolution class containing 50704 particles was refined to 5.4 Å. Particles were re-extracted using refined coordinates and unbinned (pixel size 0.889 Å/pixel) and subjected to a round of 3D refinement, Bayesian Polishing, and postprocessing/B-factor sharpening to yield a final map 4.0 Å resolution and its local resolution filtered variant calculated using Relion’s own algorithm. Local resolution maps are shown for ABCA7_BPL_ (Map1 & Map2) and ABCA7_PE_ in Figure S3.

For the ABCA7_DIGITONIN_ dataset (Figure S4B-E), a total of 7437149 particles were picked from 16213 ctf corrected and motion corrected micrographs and extracted at a 3-fold binned pixel size of 2.667 Å/ pixel. After 2D classification, 1220497 particles were subjected to 3D classification (K=8) using a low pass filtered (60 Å) map of our ABCA7_BPL_ structure as a reference. A single class comprising 324727 particles was refined to 5.4 Å. The refined coordinates were then used to re-extract unbinned particles (0.889 Å/pixel), subjected to 3D refinement, and Bayesian polishing. Further 3D classification (K=5) was performed and two similar classes containing 149590 particles were picked for 3D refinement and postprocessing/B-factor Sharpening to yield a 3.9 Å map.

For the ABCA7_EQ-ATP_ dataset, an initial set of 2660267 particles were picked from 4914 ctf corrected micrographs (Figure S5B-E). After 2D classification, 469397 particles were subjected to initial model building. This model was used as a 3D reference to perform 3D classification (k=5). A single class comprising 174415 particles was refined to 5.4 Å, the corresponding particles unbinned and re-refined. The nanodisc density was subtracted within Relion, followed by 3D classification (k=3). The highest resolution class comprising 51780 particles was refined to 4.3 Angstroms and used for 3D reference-based particle picking for a larger data set comprising the initial 4914 movies and a new set of 4474 ctf corrected micrographs. A total of 3773280 particles were picked and subjected to multiple rounds of 2D classification in Relion 4.0-beta. 898916 particles were subjected to 3D classification (K=3) and a single highest resolution class consisting of 407424 particles was refined to 5.4 Å. The refined coordinates were then used to re-extract the respective particles without binning (0.889 Å/ pixel) and refined again before 3D classification (K=5). A single class comprising 177230 particles was selected and subjected to 3D refinement, Bayesian polishing, and postprocessing/B-factor sharpening to yield a final map at 3.7 Å.

### Model building and refinement

Model building was done in coot 0.9.5^50^ using a combination of Map 1 and its local resolution filtered variant and Map 2. Both Map 1 and Map 2 displayed significant conformational heterogeneity in the second half of ABCA7, with the quality of density in Map 1 allowing placement of a TMD2 model guided in part by the homologous ABCA1 structure. Density attributed to inter-TMD phospholipids was clearest in Map 1. Map 2 revealed very poor and discontinuous density for TMD2-NBD2 but significantly better density for TMD1-NBD1 and the majority of ECD1 and ECD2, allowing for de novo model building. The model for ECD was also guided by the presence of nine glycosylation sites (N78, N98, N312, N340, N1335, N1381, N1386, N1457, & N1518) as well as 4 disulfide bond pairs. Density for the lid region of the ECD was missing in both maps. Model building for both NBDs was guided by structures of the homologous transporters TM287^51^, ABCG2, and ABCA1, where density features did not allow for de novo model building. We observed extra density at the nucleotide binding sites for both NBDs despite their open conformation. The structure of the RD was based on a homology model of the predicted RD structure in ABCA4^25^. The quality of density for the ABCA7 RD allowed rigid body placement for the entire domain. Restrained real space refinement of the model was carried out in Phenix 1.19.1^52^ using automatically generated secondary structure restraints. Structural figures were prepared in UCSF Chimera v. 1.13.1^53^, ChimeraX v. 1.2.5^54^, and PyMOL 2.4.1 (The PyMOL Molecular Graphics System, Version 1.8 Schrödinger, LLC).

The model for ABCA7_PE_ was generated by rigid body placement of the ABCA7_BPL_ model followed by real space refinement in phenix as described above. The model for ABCA7_DIGITONIN_ was generated by rigid body fitting each TMD, NBD, ECD, and RD into its postprocessed map followed my manual adjustment of sidechains as allowed for the map. Non proteinaceous EM density was modeled as a CHS and Digitonin (ligand ID Y01) molecule. The corrected model was real space refined against the postprocessed map. The model for ABCA7_EQ-ATP_ was built starting with a homology model based on the ATP bound structure of ABCA4 (PDB 7LKZ), followed by manual adjustment of the structure as required and permitted by the map. The ECD was replaced by a rigid body fitted model of the ECD from the ABAC7_Digitonin_ structure. This model was then refined against both the postprocessed map, and its local resolution filtered counterpart.

### MD simulations

We employed MD simulations to capture the arrangement and dynamics of the lipid bilayer induced by the experimentally derived open conformation of ABCA7_PE_, which was used as the starting model in all the simulations. For system setup, a C-terminal carboxylate capping group, an N-terminal ammonium capping group and all the hydrogen atoms were added using the Psfgen plugin of VMD (Visual Molecular Dynamics)^56^. The resulting all-atom (AA) model was then converted to a coarse-grained (CG) Martini model using the Martinize protocol (http://www.cgmartini.nl/), using an elastic network on atom pairs within a 10-Å cutoff. The Orientations of Proteins in Membranes (OPM) database^57^ was used to identify and align the transmembrane region of the protein with the membrane normal. The protein was embedded in two distinct lipid bilayers (palmitoyl-oleoyl-phosphatidyl-ethanolamine (POPE) and cholesterol with a molar ratio of 4:1 (POPE/Chol), and palmitoyl-oleoyl-phosphatidyl-choline (POPC) and cholesterol with a molar ratio of 4:1 (POPC/Chol), respectively). The protein secondary structure was defined from the AA model and was maintained throughout the CG simulations by the applied elastic network. To increase the sampling of lipid-protein interactions and improve statistics, four independent copies of the CG protein were placed at a distance of 200 Å in a large lipid bilayer (400×400 Å^2^). The system was then solvated and ionized with 150 mM salt using Insane^58^.

The systems were simulated employing GROMACS 2021.3^59,60^. A 20-fs timestep was employed in all the simulations. The temperature was maintained at 310 K with a velocity-rescaling thermostat^61^ employing a coupling time constant of 1 ps. A semi-isotropic 1 bar pressure was maintained using the Berendsen barostat ^62^ with a compressibility and relaxation time constant of 3×10^-4^ bar and 5 ps, respectively. The systems were energy minimized for 1,000 steps, followed by short equilibration runs of 18 ns, while restraints were applied to lipid bilayer headgroups and protein backbones. During this time the restraints on bilayer headgroups were reduced gradually from k = 200 kJ.mol^-1^.nm^-2^ to zero, whereas the protein backbones’ restraints (k = 1,000 kJ.mol^-1^.nm^-2^) were kept constant. Each system was then simulated for 2 μs, with restraints only applied to the protein backbones, resulting in an aggregate sampling of 8 μs (4 copies × 2 μs). All the systems were simulated following the same MD protocol.

All the molecular images were generated using VMD^56^. The membrane deformation induced by ABCA7 was quantified by calculating the *z* distance of the lipid phosphate moieties (PO_4_ bead type in Martini) with respect to the bilayer midplane, over the last 1 μs of each trajectory. The generated histogram (binned in 2×2 Å^2^ bins) in each leaflet illustrates the spatial distribution of the height of the lipid head groups within each leaflet. We quantified the differential movement of POPE and POPC within the protein lumen by calculating the number of phospholipids located within the TMDs. If the PO_4_ bead of a phospholipid was within 22.5 Å and 12.5 Å in *x* and *y*, respectively, with respect to a protein’s center in the *x-y* plane (membrane plane), then the phospholipid was considered to be within the TMD lumen (Figure S9C).

## Acknowledgments

We would like to thank Dr. Kaspar Locher at ETH, Zurich, Switzerland, for providing the synthetic gene construct of ABCA7. We would also like to thank the cryo-EM and shared instruments core facilities at the Hormel Institute for help with experimental setup, and Dr. Rhoderick Brown, Dr. Jarrod French and Dr. Jeppe Olsen for critical reading and discussion during manuscript preparation. This work was supported in part by the Hormel Foundation (Institutional research funds to AA), the National Institutes of Health (NIH) award 1R21-AG069180-01A1 (to AA), and the Cure Alzheimer’s fund (to TK). The computational component of the project was supported by the NIH awards P41-GM104601 (to ET) and R01-GM123455 (to ET). We also acknowledge computing resources provided by Blue Waters at National Center for Supercomputing Applications (NCSA), and by eXtreme Science and Engineering Discovery Environment (XSEDE) (grant MCA06N060 to ET), and by Microsoft Azure.

## Author Contributions

AA conceived the research. LTML, JRT, and AA performed all experiments. MD simulations were conducted and analyzed by SD, SP, and ET. TK participated in assay design. LTML, JRT, and AA wrote the manuscript with input from all other authors.

## Data and materials availability

The cryo-EM Maps have been deposited at the Electron Microscopy Databank (EMDB) under accession codes EMD-AAAAA (ABCA7_BPL_ Map 1), EMD-BBBBB (ABCA7BPL Map 2), EMD-CCCCC (ABCA7_PE_), EMD-DDDDD (ABCA7_EQ-ATP_), and EMD-EEEEE (ABCA7DI_GITONIN_). The associated atomic coordinates have been deposited at the Protein Data bank (PDB) under accession codes 1WWW (ABCA7_BPL_), 2XXX(ABCA7_PE_), 3YYY, (ABCA7_EQ-ATP_), and 4ZZZ (ABCA7DI_GITONIN_).

---

**Figure S1.**
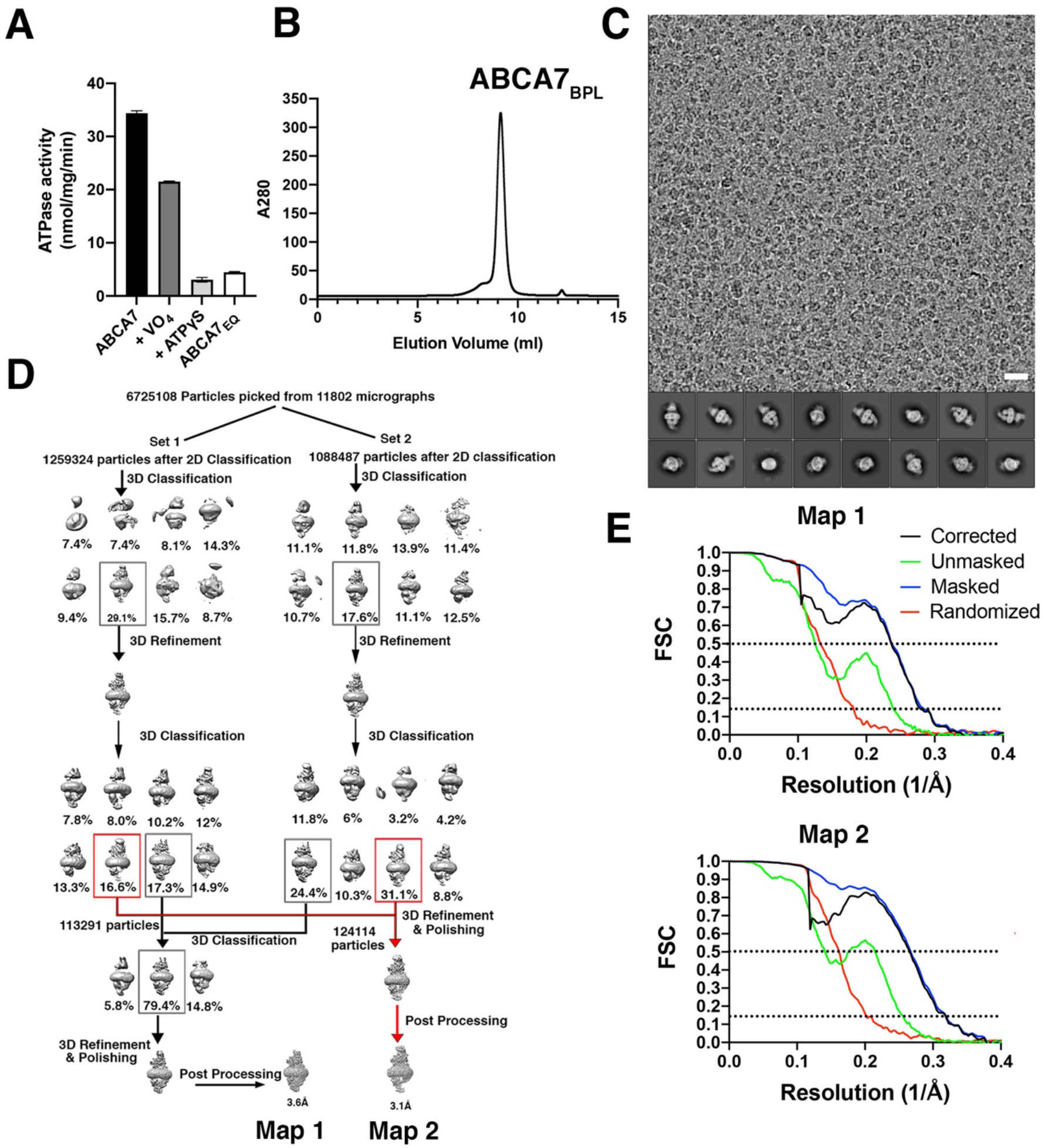
ABCA7_BPL_ characterization and Cryo-EM data processing. **A** ATPase data for nanodisc reconstituted ABCA7 with and without sodium orthovanadate (VO4) or ATPγS. N=3 and error bars represent s.d **B** SEC peak of ABCA7_BPL_ sample for cryo-EM. **C** Representative cryo-EM micrograph at -2.5 μm defocus. Scale bar = 20 nm. **D** cryo-EM processing workflow. Boxes indicate 3D classes used for further refinement for both Map 1 and Map 2 (red). **E** Fourier shell correlation (FSC) curves for Map 1 (top) and Map 2 (bottom) Dotted lines indicate position 0. 143 and 0.5 cutoff criteria for resolution estimates.

**Figure S2.**
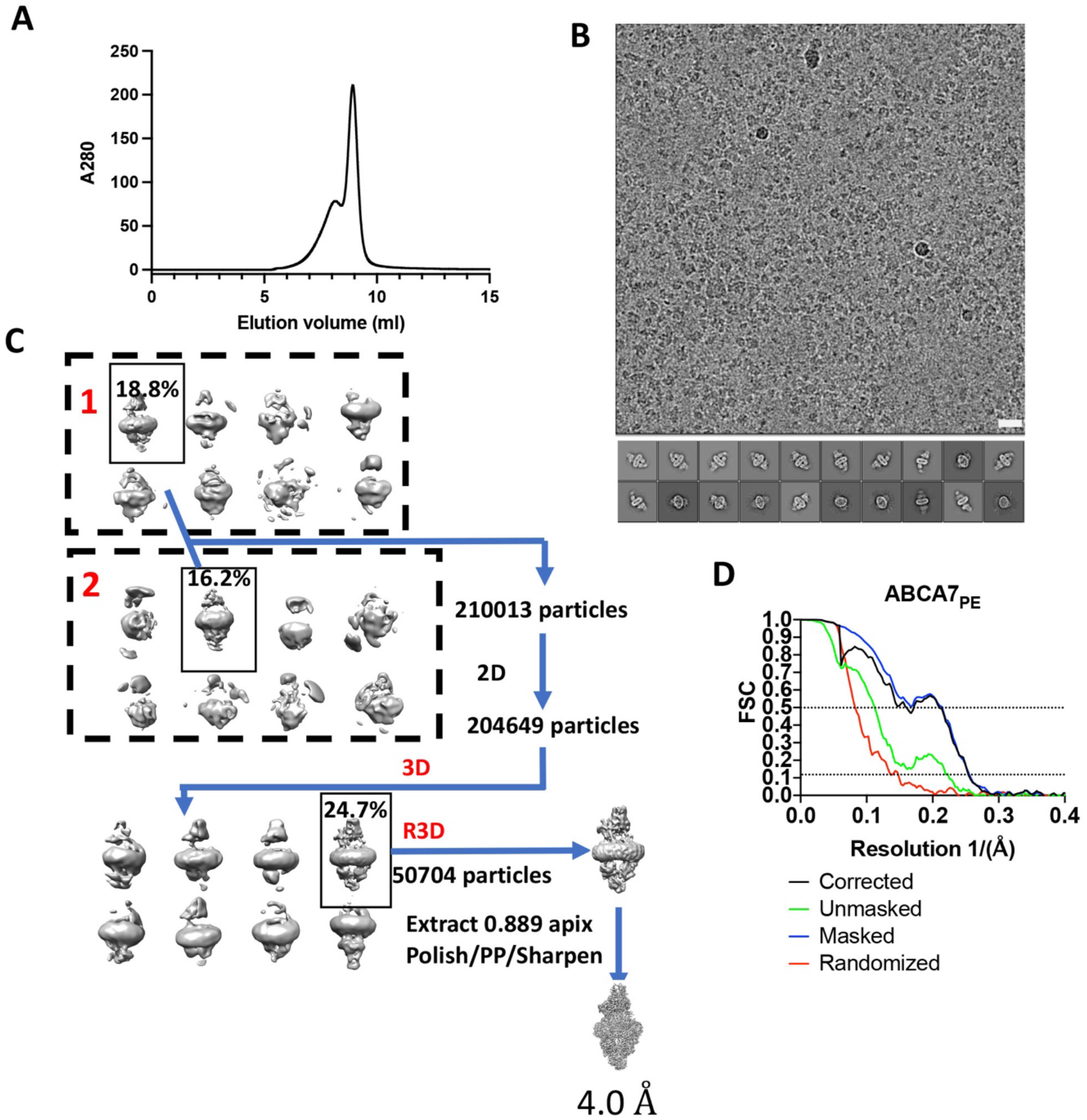
ABCA7_PE_ Cryo-EM data processing. **A** Size exclusion chromatography micrograph of cryo-EM sample showing monodisperse ABCA7_PE_ nanodisc (main peak). **B** Representative micrograph at -2.5 μm defocus and 2D classes. Scale bar = 20 nm. **C** cryo-EM processing workflow. Dashed boxes demarcate Subsets 1 and 2. Solid boxes indicate 3D classes used for further refinement. C2D = 2D Classification, C3D=3D classification, R3D = 3D refinement **D** Fourier shell correlation (FSC) curves for ABCA7_PE_ Dotted lines indicate position 0.143 and 0.5 cutoff criteria for resolution estimates.

**Figure S3.**
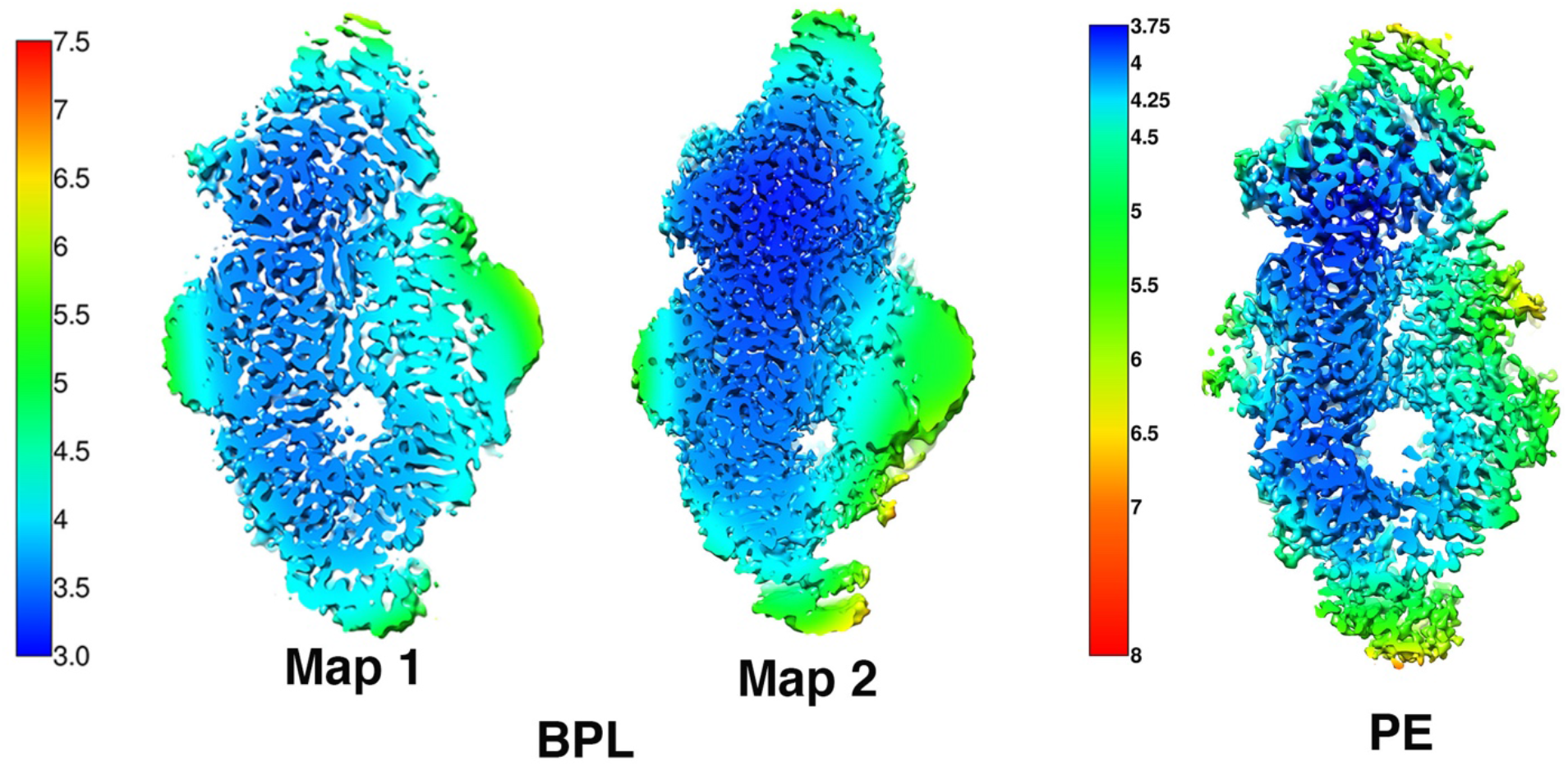
ABCA7_BPL_ and ABCA7_PE_ local resolution filtered maps. Color keys are indicated on the left of each set of maps with numbers representing resolution (Å).

**Figure S4.**
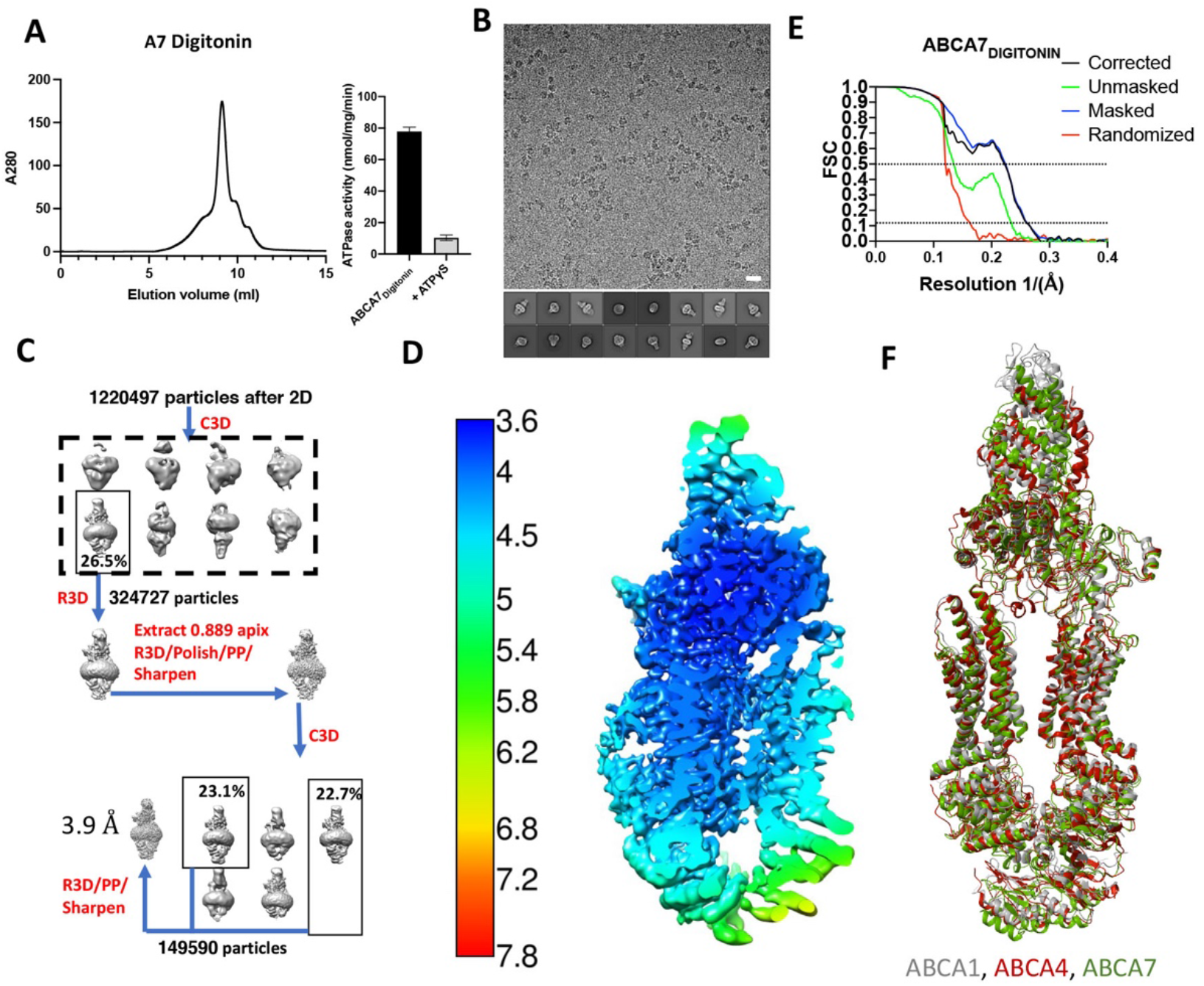
ABCA7DIGITON cryo-EM processing. **A** SEC profile of ABCA_7DIGITONIN_ and its ATPase activity with and without ATPγS. **B** Representative micrograph at -2.5 μm defocus and 2D classes. Scale bar = 20 nm. **C** cryo**-**EM processing workflow. C2D = 2D Classification, C3D=3D classification, R3D = 3D refinement. **D** Local resolution colored EM map of ABCA7_DIGITONIN._ **E** Fourier shell correlation (FSC) curves for ABCA7_DIGITONIN._ Dotted lines indicate position 0.143 and 0.5 cutoff criteria for resolution estimates. **F** Superposition of ABCA7 (green, this manuscript), ABCA1 (grey, PDB 5XJY), and ABCA4 (red, PDB 7LKP) structures in digitonin.

**Figure S5.**
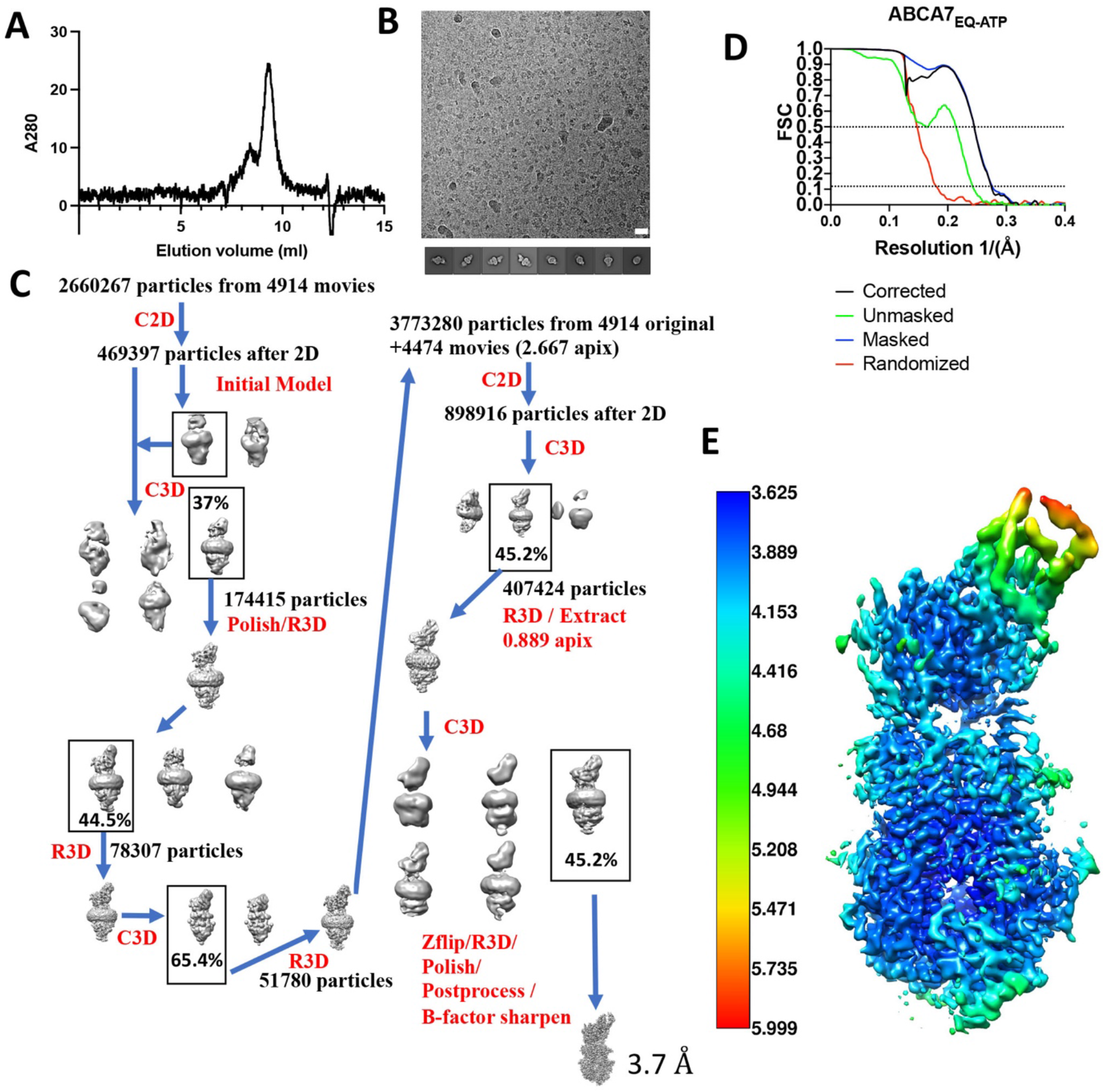
ABCA7_EQ-ATP_ cryo-EM processing. **(A)** SEC peak for ABCA7_EQ-ATP_ in nanodiscs. **B** Representative EM micrograph (−2.5 defocus) and rep 2D classes for ABCA7_EQ-ATP_ sample. **C** Cryo-EM data processing pipeline. C2D = 2D Classification, C3D=3D classification, R3D = 3D refinement. **D** FSC curves. **E** Local resolution colored EM map.

**Figure S6.**
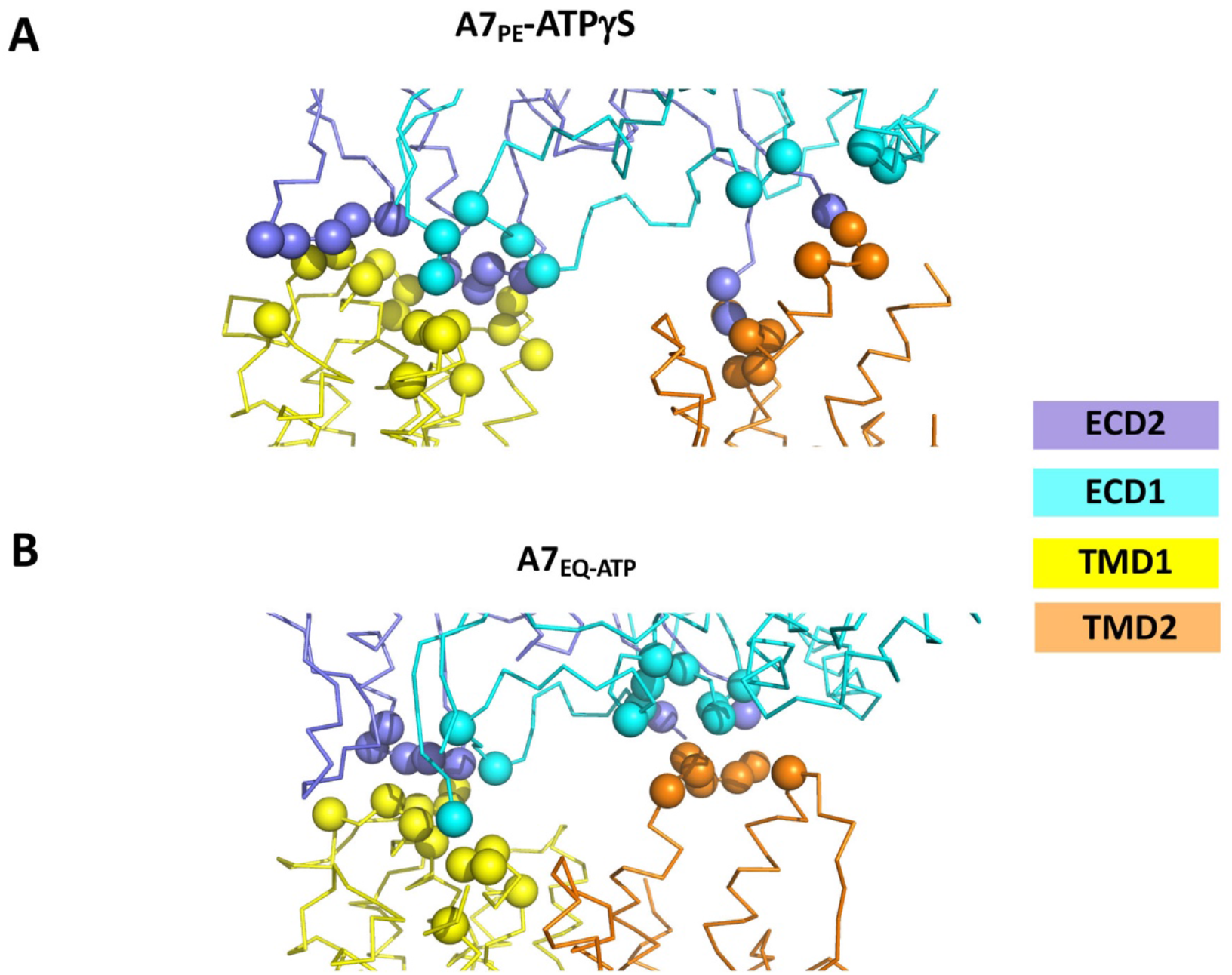
TMD-ECD interfaces of open and closed form ABCA7. **(A)** The TMD-ECD binding interfaces of ABCA7_PE_ with TMD1 and TMD2 and with Cα for residues in TMD1 and TMD2 within 5Å of either ECD or vice versa shown as spheres. **(B)** The same analysis for the TMD-ECD binding interfaces of ABCA7_EQ-ATP_ with closed cavity.

**Figure S7.**
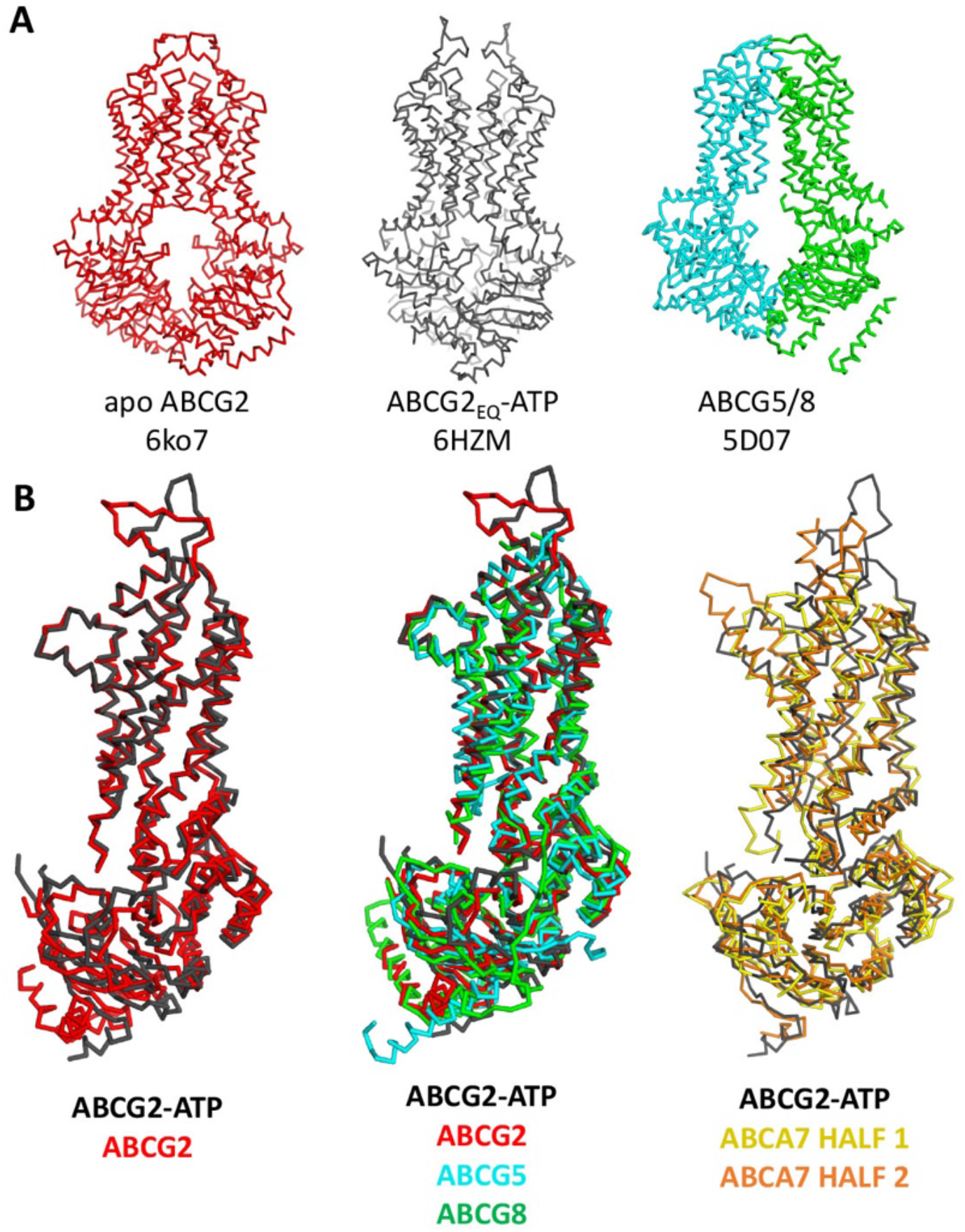
Conservation of ABCA and ABCG family structural elements. **(A)** Ribbon representation of select ABCG family transporter structures and their respective PDB IDs including apo open ABCG2 (red), closed ATP bound structure of ABCG2_EQ_ (black), and ABCG5/G8 (cyan and green, respectively) (**B**) Alignment of NBD-TMD pairs from open and closed conformations of ABCG2 (left), ABCG2 and ABCG5/G8 (center), and closed ABCG2 with TMD-NBD pairs from ABCA7 half 1 (gold) and half 2 (orange).

**Figure S8.**
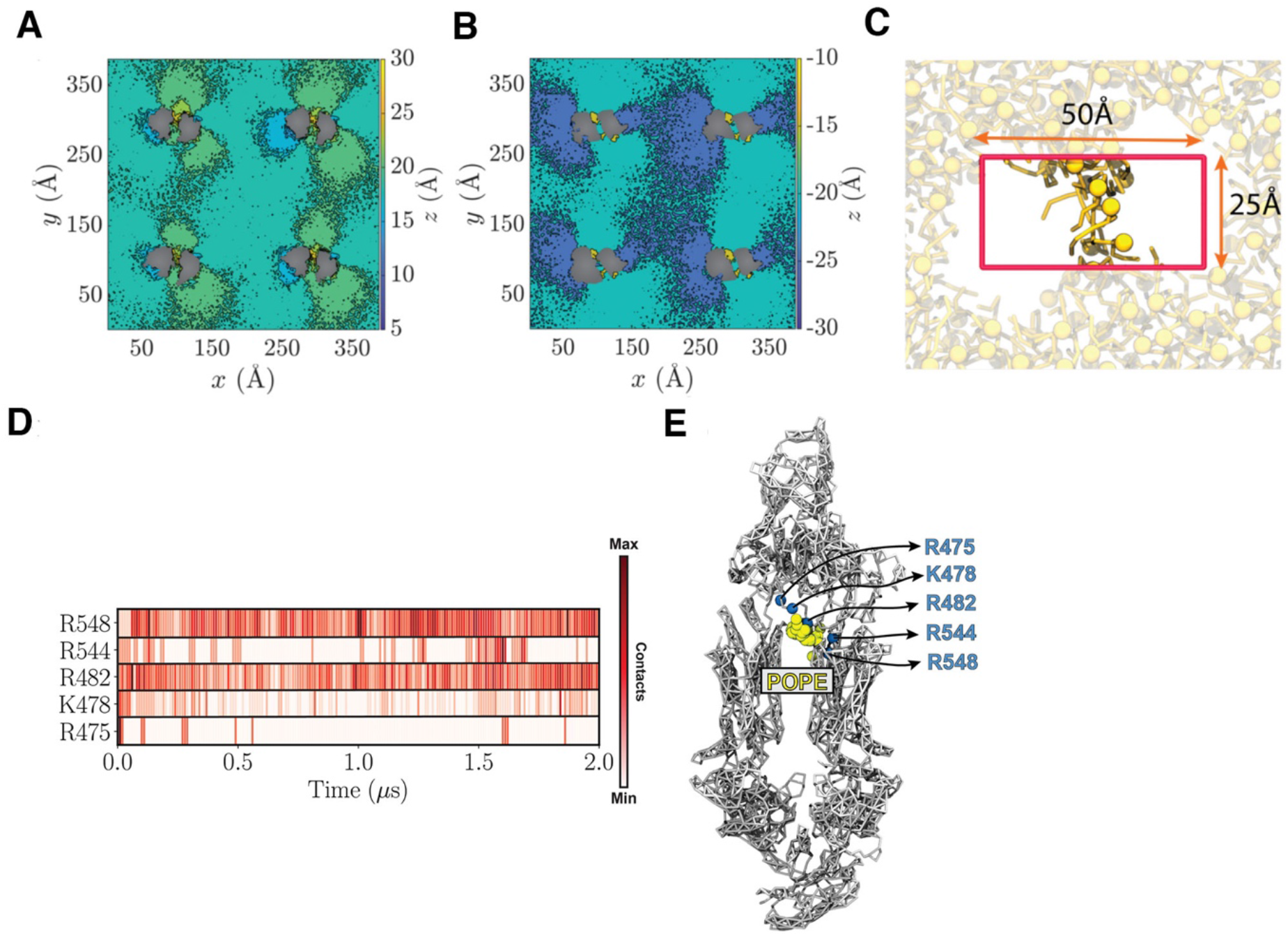
Lipid configuration induced by protein copies in a POPC bilayer and criteria used for identifying luminal lipids in MD simulations. **(A/B)** POPC headgroup height calculated for extracellular (A) and cytoplasmic (B) leaflets. **(C)** Dimension and position of the box used to calculate the number of phospholipids (yellow) partitioned in the TMD leaflets. The box is centered at the protein center and has dimensions of 50 Å and 25 Å in *x* and *y* directions, respectively. The ABCA7 transporter is hidden for clarity. **(D)** Contact map, based on the number of POPE headgroups in proximity of each residue throughout the simulation. **(E)** Accumulation of POPE headgroups (yellow spheres) in close proximity of most frequently contacted residues throughout the simulation.

**Figure S9.**
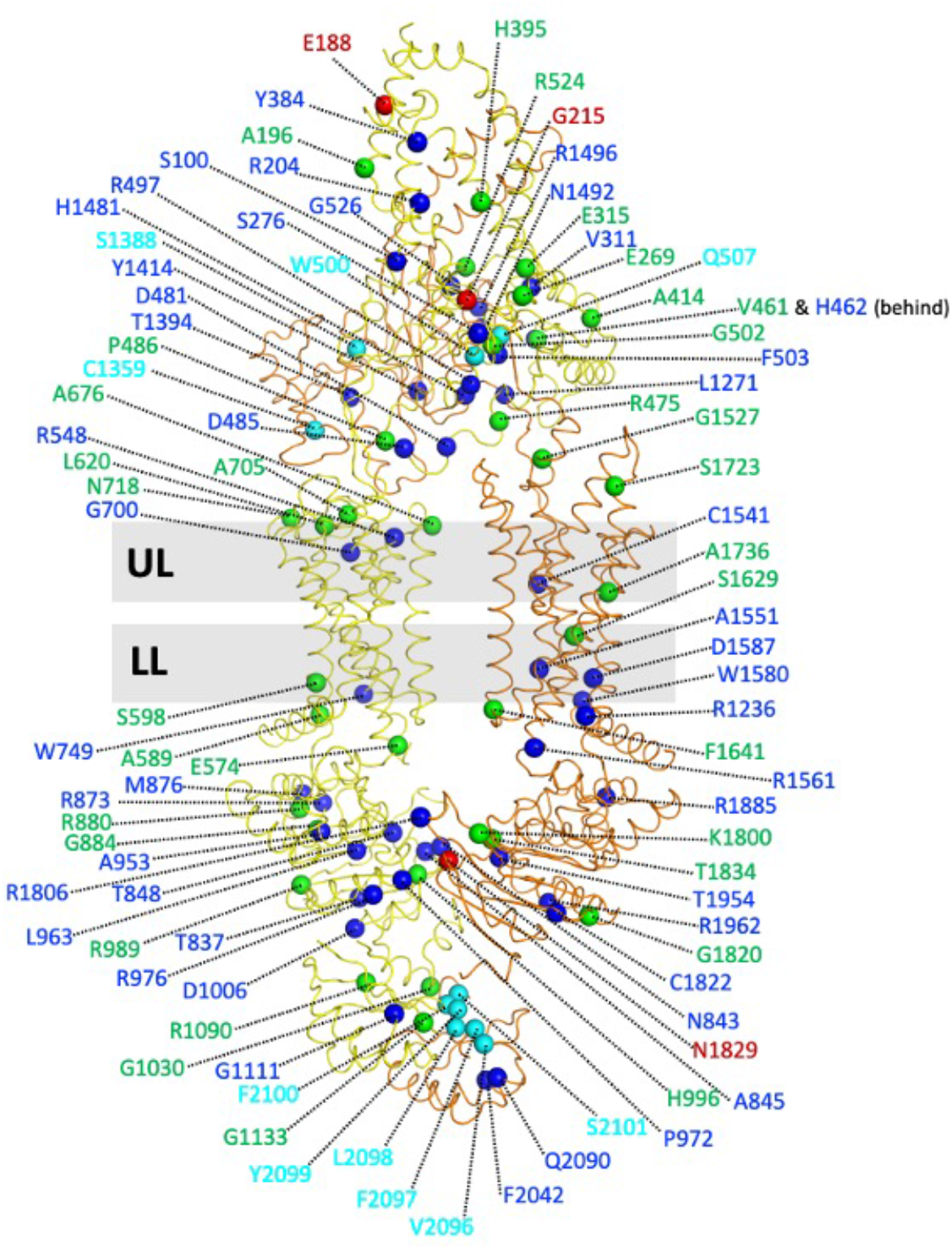
Amino acid positions of ABCA7 with variants associated with risk of AD. Structure of ABCA7 with Cα atoms shown for amino acid with known pathogenic variants (green), protective variants (red), and ABCA7 residues conserved in ABCA1 with mutations known to disrupt the latter’s binding to apoA1 and/or phospholipid translocation (blue). Cyan spheres highlight residue positions of ABCA1 mutations known to disrupt phospholipid efflux including those equivalent to the VFVNFA motif within the ABCA1 RD. Grey bars indicate the upper (extracellular) and lower (cytoplasmic) membrane bilayer leaflets (UL and LL, respectively).

**Figure S10.**
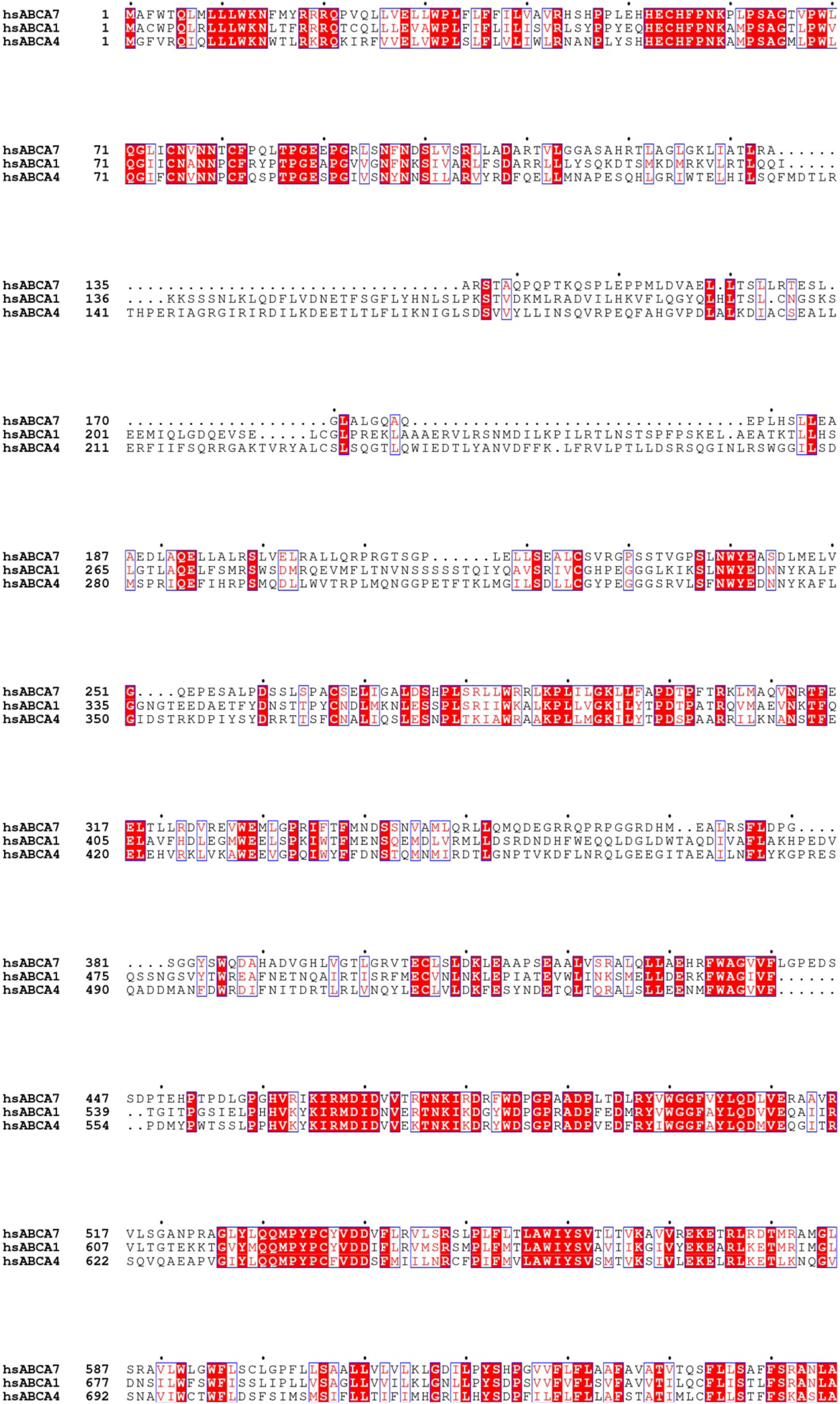
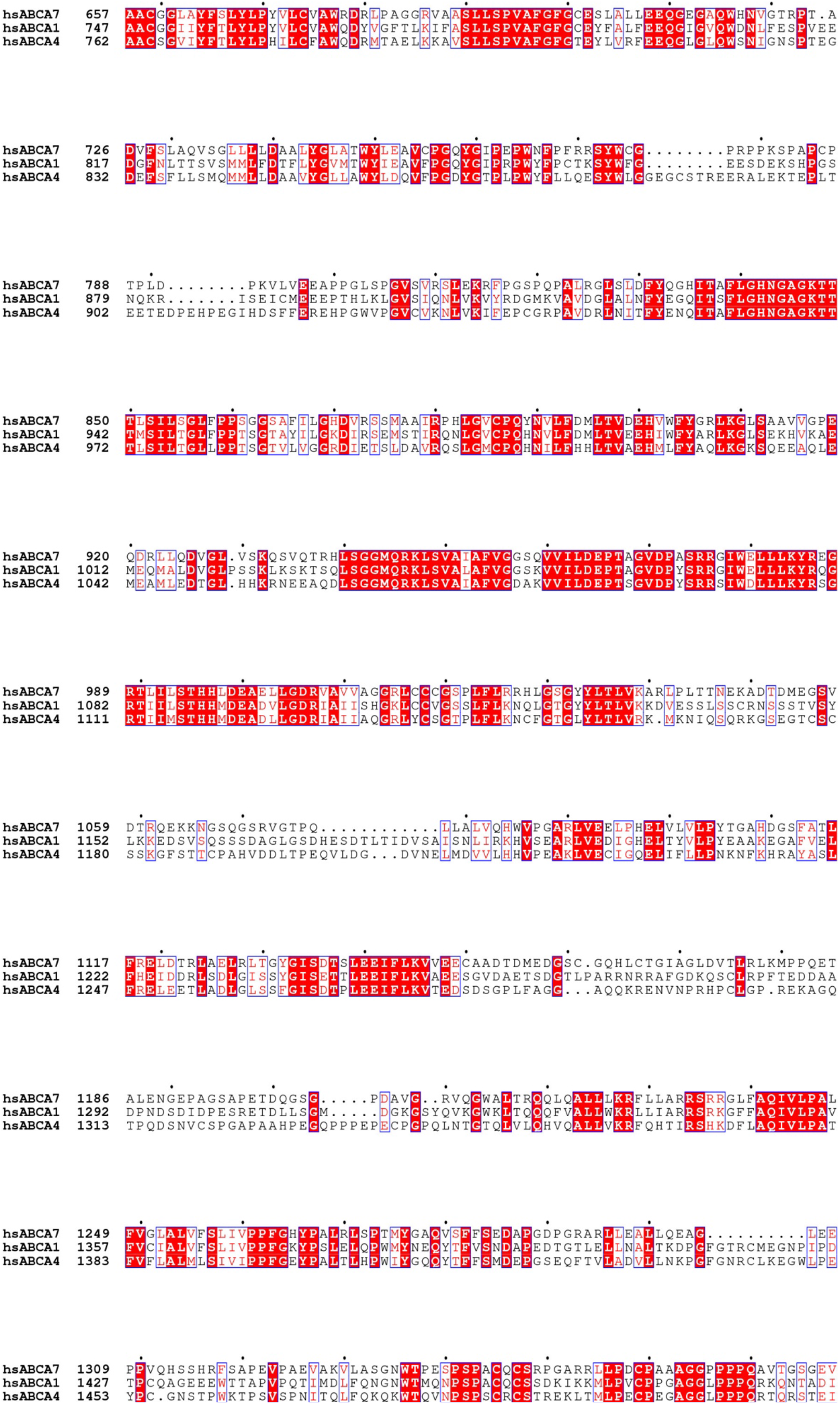
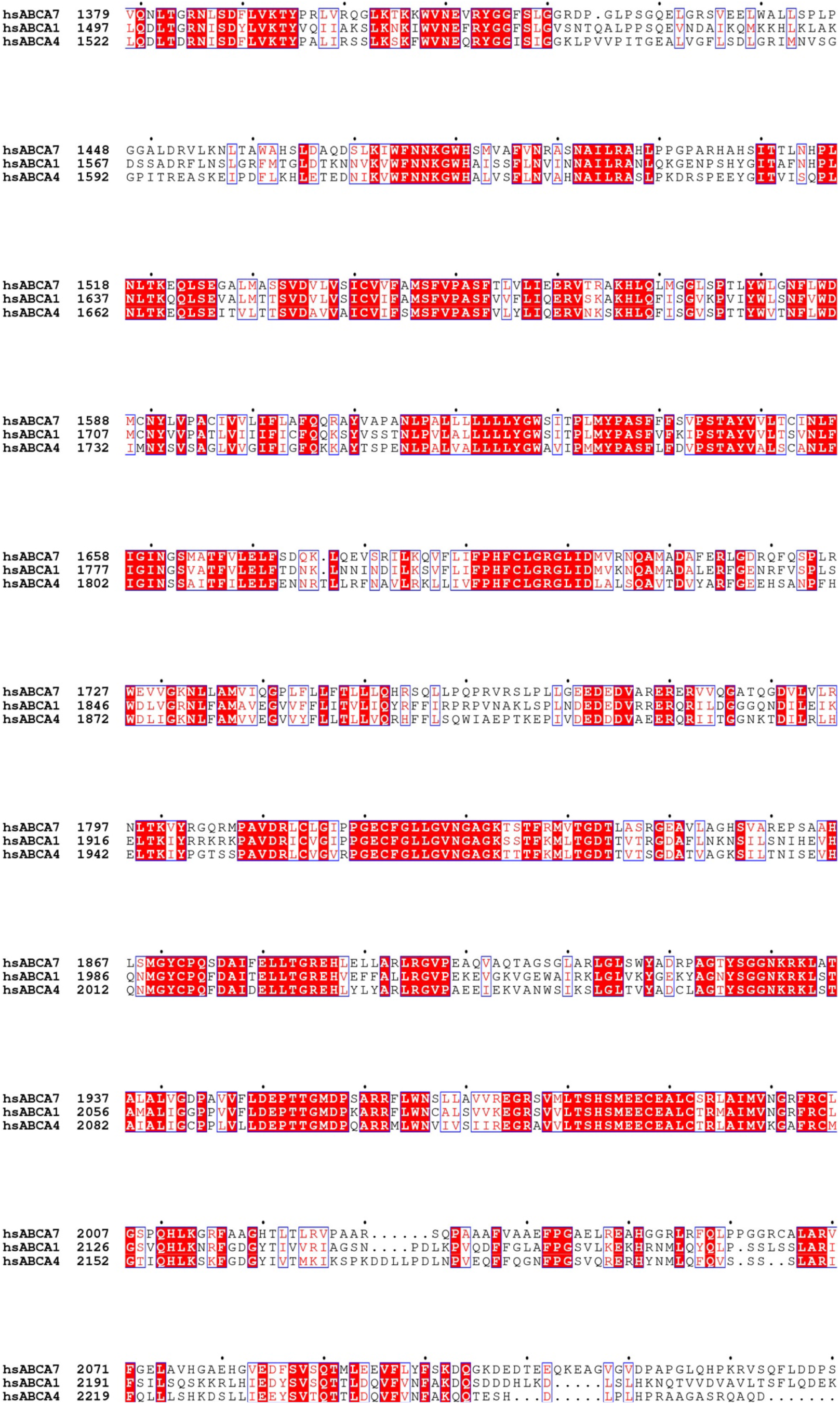
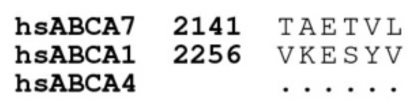
Sequence alignment of human (hs) ABCA7, ABCA1, and ABCA4.

**Table S1.** Data collection and refinement statistics.

|  |  |  |  |  |  |
| --- | --- | --- | --- | --- | --- |
| Dataset | ABCA7<sub>BPL</sub> |  | ABCA7<sub>PE</sub> | ABCA7<sub>DIGITONIN</sub> | ABCA7<sub>EQ-ATP</sub> |
| Magnification | 96k |  | 96k | 96k | 96k |
| Pixel Size (Å) | 0.895 |  | 0.889 | 0.889 | 0/889 |
| Total Dose (e/Å<sup>2</sup>) | 60 |  | 40 | 40 | 40 |
| Defocus Range (um) | -0.8 to 2.6 |  | -0.8 to 2.6 | -0.8 to 2.6 | -0.8 to 2.6 |
| Maps | Map 1 | Map 2 | Map3 | Map4 | Map5 |
| EMDB ID | EMD-22996 | EMD-22998 | EMD-A | EMD-B | EMD-C |
| # Particles in final Class | 91381 | 124114 | 50704 | 149590 | 177230 |
| Resolution (Å) (0.143 threshold) | 3.6 | 3.2 | 4.0 | 3.9 | 3.7 |
| Sharpening B factor |  |  |  |  |  |
| **Refined Coordinates** | ABCA7<sub>BPL</sub> |  | ABCA7<sub>PE</sub> | ABCA7<sub>DIGITONIN</sub> | ABCA7<sub>EQ-ATP</sub> |
| PDB ID | AAAA |  | BBBB | CCCC | DDDD |
| # Residues/Non-hydrogen Atoms | 1801/14761 |  | 1846/14749 | 1856/4662 | 1871/14748 |
| Glycans | 16 |  | 16 | 26 | 16 |
| Ligands | 21 |  | 20 |  | 2 |
| R.M.S deviations |  |  |  |  |  |
| Bond Length (Å) | 0.003 |  | 0.003 | 0.003 | 0.003 |
| Bond Angles (°) | 0.616 |  | 0.600 | 0.678 | 0.645 |
| MolProbity Statistics |  |  |  |  |  |
| MolProbity Score | 1.68 |  | 1.72 | 1.83 | 1.77 |
| Clashscore | 8.05 |  | 8.16 | 11.40 | 9.83 |
| Poor rotamers (%) | 0.00 |  | 0.00 | 0.00 | 0.00 |
| Ramachandran statistics |  |  |  |  |  |
| Favored (%) | 96.38 |  | 95.84 | 96.24 | 96.22 |
| Allowed (%) | 3.62 |  | 4.16 | 3.76 | 3.78 |
| Outliers (%) | 0.00 |  | 0.00 | 0.00 | 0.00 |

